# The evolution of gene functional repertoire in Amorphea: Divergent strategies across Amoebozoa, Fungi and Metazoa

**DOI:** 10.1101/2025.05.13.653667

**Authors:** Alex Gàlvez-Morante, Cédric Berney, Daniel J. Richter

**Affiliations:** Institut de Biologia Evolutiva (CSIC-Universitat Pompeu Fabra), Passeig Marítim de la Barceloneta 37-49, 08003 Barcelona, Spain

**Keywords:** Amoebozoa, Metazoa, Fungi, evolution, Amorphea, ancestral reconstruction, COG, Pfam

## Abstract

Metazoa and Fungi have been extensively studied to reconstruct the trajectory of Opisthokont evolution. Their sister group, Amoebozoa, provides additional potential to generate valuable insights into the origins of Opisthokont lineages. Amoebozoa represent a diverse group of amoeboid organisms, which have adapted to a wide range of environments and ecological niches. Studying Amoebozoa not only helps to illuminate Opisthokont evolution but also reveals the mechanisms that have driven their ecological success.

Here we report the discovery of *Apostamoeba explorator* strain BEAP0066, representing a novel lineage within Amoebozoa with intriguing behaviors like the “double-amoeba”, a behavior characterized by the bipolarization of a cell into two poles that coexist and act as two semi-independent cells; and the “colonizing rings”, the generation of a front of amoebae advancing together and grazing on bacterial mats.

By analyzing the gene content of *A. explorator* and diverse amoebozoans with ancestral gene content reconstructions, correspondence analyses of Clusters of Orthologous Groups (COG) category composition and Pfam clan clustering, we revealed distinct evolutionary trajectories for Amoebozoa, Metazoa, and Fungi. Amoebozoa retained an ancestral Amorphea-like state, characterized by an enrichment of genes related to motility, phagocytosis, and environmental adaptability; while Metazoa specialized in multicellularity-related genes and Fungi in metabolism and transport. These findings suggest that retention of gene function composition, rather than gene loss, played a key role in shaping Amoebozoa evolution.

**SIGNIFICANCE STATEMENT:** Amoebae are cells with no defined shape that move by extending temporary cell membrane projections, usually composed of actin, called pseudopodia. Amoeboid cells are present in all major eukaryotic lineages, from the well-known *Amoeba proteus* in Amoebozoa to human macrophages in Metazoa, *Salpingoeca rosetta* in Choanoflagellatea, *Vampyrella lateritia* in Rhizaria, and other examples such as *Naegleria fowleri* in Heterolobosea. The prevalence of this lifestyle highlights the evolutionary success of the amoeboid form and its ability to adapt to diverse environmental conditions and ecological roles. The study of amoebae and amoeboid cell types is crucial for advancing research in medicine, ecology, and evolution. Understanding the phylogeny of Amoebozoa remains a key focus in phylogenomics, as deep divergences within the group complicate the taxonomic placement of certain taxa and have implications for character evolution. Thus, the discovery, identification, and characterization of new Amoebozoa species can help resolve current uncertainties. Recent studies have identified divergent trajectories in gene content composition within one lineage of Amorphea, the Opisthokonts. Fungi evolved through an expansion of metabolic genes, whereas Metazoa accumulated genes associated with multicellularity. Inspired by these findings, we analyzed the proteomes of Amoebozoa, Metazoa, and Fungi. Using correspondence analysis of the relative composition of Clusters of Orthologous Groups (COG), ancestral reconstruction and the analysis of Pfam domain clan presence in supergroup-specific gene clusters, we aimed to determine whether the three supergroups within Amorphea exhibit distinct clustering patterns. Our results provide evidence of divergent functional evolution in Amoebozoa, Fungi, and Metazoa. Amoebozoa retained an ancestral Amorphea-like state, characterized by an enrichment of genes related to motility, phagocytosis, and environmental adaptability; while Metazoa specialized in multicellularity-related genes and Fungi in metabolism and transport. These findings suggest that retention of gene function composition, rather than gene loss, played a key role in shaping Amoebozoa evolution.

## INTRODUCTION

Amoebae are cells with no defined shape that move through the extension of temporary cell membrane projections, usually composed by actin, called pseudopodia. Amoeboid cells are present in all major eukaryotic lineages (Brunet et al., 2021), from the well-known *Amoeba proteus* in Amoebozoa (Pomorski et al., 2007) to human macrophages in Metazoa and *Salpingoeca rosetta* in Choanoflagellatea (Brunet et al., 2021), *Vampyrella lateritia* in Rhizaria (Hülsmann & Grębecki, 1995) and many other examples like *Naegleria fowleri* in Heterolobosea (Siddiqui et al., 2016) or *Amphitrema stenostoma* in Stramenopiles (González Miguéns, 2023). The prevalence of this lifestyle is evidence of the evolutionary success of the amoeboid form and its ability to adapt to different environmental conditions and ecological roles.

The study of amoeba and amoeboid cell types is crucial in order to shed light to different fields like medicine, as they are a part of our immune system (Park et al., 2022), microbiome (Stensvold et al., 2023) and pathogens (Wang et al., 2023); ecology, as amoeba occupy a wide variety of ecological niches, acting as predators (Hess & Suthaus, 2022), producers (Gabr et al., 2022) and parasites (Borkens, 2024); or evolution, as amoeboid cells are widely distributed among the different eukaryotic lineages. The single largest amoeboid taxon, Amoebozoa, is the closest known relative of Obazoa, the group containing both Fungi and Metazoa (Tekle et al., 2022).

Amoebozoans main cell type is the trophozoite, the active and feeding stage of the amoeba, but they are able to undergo several different life cycle transitions. One widespread example would be the cyst, a dormant life stage generated to resist harsh environmental conditions, but many are able to generate other life stages like flagellates and spores, among other reproductive and social behaviors (Schilde & Schaap, 2013). Understanding the phylogeny of Amoebozoa is an active topic in the phylogenomics field, as deep divergences make it difficult to produce robust taxonomic placement of some key taxa, with implications for character evolution in the group (Pawlowski & Burki, 2009). Thus, the discovery, identification and characterization of new Amoebozoa species can contribute to resolving current uncertainties.

In this project, we isolated and cultured a new and puzzling amoeba BEAP0066 that complements extant phylogenetic studies, due to its novel phylogenetic placement and divergence from currently described groups. We sequenced the transcriptome of BEAP0066 and observed numerous behaviors and life history stages via time-lapse microscopy.

Recently, divergent trajectories in gene content composition were identified in a comparison of the evolution of Fungi, which was characterized by an expansion of metabolic genes, and Metazoa, which accumulated genes potentially important for their multicellularity (Ocaña-Pallarès et al., 2022). These findings inspired us to use BEAP0066’s predicted proteome, alongside a deep sampling of Amoebozoa, Metazoa and Fungi, to perform correspondence analysis of the relative composition of Clusters of Orthologous Groups (COG), replicating a similar methodology to the proposed in Ocaña-Pallarès *et al*. The goal was to determine whether the three studied supergroups within Amorphea exhibit independent clustering patterns, which was indeed observed in our results, providing evidence of divergent evolution in Amoebozoa as well as Fungi and Metazoa. Additionally, we conducted an ancestral reconstruction of gene family evolution across Amorphea to compare the COG profiles of ancestral species with those of extant supergroups. To complement these analyses, we identified overrepresented Pfam domain clans in gene clusters specific per Amoebozoa, Fungi and Metazoa to better understand the changes in functional potential that drove the evolution of Amorphea.

## RESULTS

### Phylogenetic placement of BEAP0066

BEAP0066’s 18S small subunit ribosomal RNA gene sequence is highly divergent and could not be included in any known clade (Supplementary Figure 1). To elucidate the phylogenetic position of BEAP0066 within Amoebozoa, we sequenced its transcriptome and reconstructed a phylogenomic tree with PhyloFisher (Tice et al., 2021) followed by IQ-TREE (Nguyen et al., 2015) (Figure 1). Our new isolate branches inside Discosea, between Himatismenida and the lineage containing Pellitida and Acanthopodida. The high divergence of BEAP0066 coupled with its phylogenetic position as sister group to Pellitida and Acanthopodida, while Himatismenida is sister to the clade containing Pellitida + Acanthopodida + BEAP0066, makes it problematic to assign BEAP0066 to a known Amoebozoa order.

**Figure 1.**
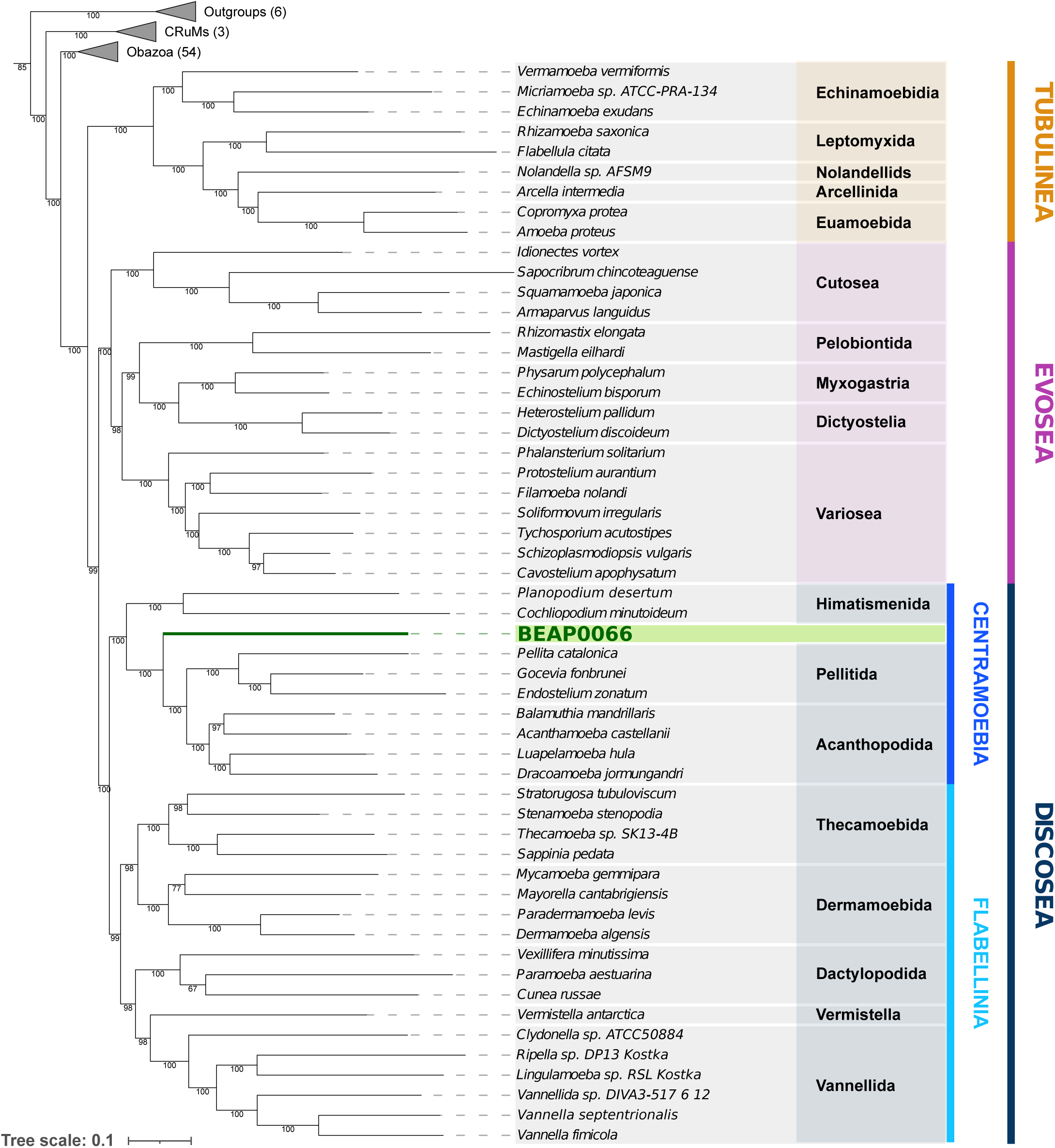
BEAP0066 and Vannella septentrionalis phylogenetic position within Amoebozoa. Phylogenetic tree of Amoebozoa diversity based on 230 genes, with new sequences from BEAP0066 and V. septentrionalis, reconstructed with Phylofisher and IQ-TREE using the LG+R10 model of sequence evolution. Node support reflects 1000 bootstraps. BEAP0066 branches with full support within Discosea as a sister to Pellitida + Acanthopodida. V. septentrionalis branches as sister to Vannella firmicola, as expected. Number of species included in each outgroup is indicated in parenthesis. See Supplementary file 1 for the phylogenetic tree with all outgroup species represented. The tree was manually rooted between Amorphea + CRuMs and Outgroups (containing Malawimonadidae, Diaphoretickes and Ancyromonadida).

In addition, for this project, we sequenced the transcriptome of another amoeba from within Discosea, whose 18S sequence and morphology match *Vannella septentrionalis*. We included *V. septentrionalis* in the phylogenomic tree that we reconstructed, where we recovered an expected placement inside Vannellida.

### Morphology and behavior of BEAP0066

We identified numerous cell type morphologies in BEAP0066 cell culture (Figure 2, Supplementary Figure 2). BEAP0066 trophozoites (Figure 2F) present a variable cell length, which (excluding pseudopodia) ranges between 4 - 17 μm (average 7.9 μm, n = 20); the cell breadth is also variable and ranges from 2 - 7 μm (average 2.3 μm, n = 20); and the length/breadth ratio (L/B) varies from 1 - 5 (average 2.3, n = 20). The variability in these measurements depended on parameters such as growth temperature, nutrient concentration or density of prey bacteria. Consistent with the morphology of other members of Discosea, the locomotive stage of the cell is flat and shows multiple filopodia of variable size that can extend several cell lengths, in contrast to the filopodia of other Acanthapodida species. The size of the nuclei ranges between 1 - 2.5 μm (average 1.7, n = 20). Multinucleated cells can often be found in the culture (e.g., Figure 2G,H), and their frequency is higher in cells of a larger size.

**Figure 2.**
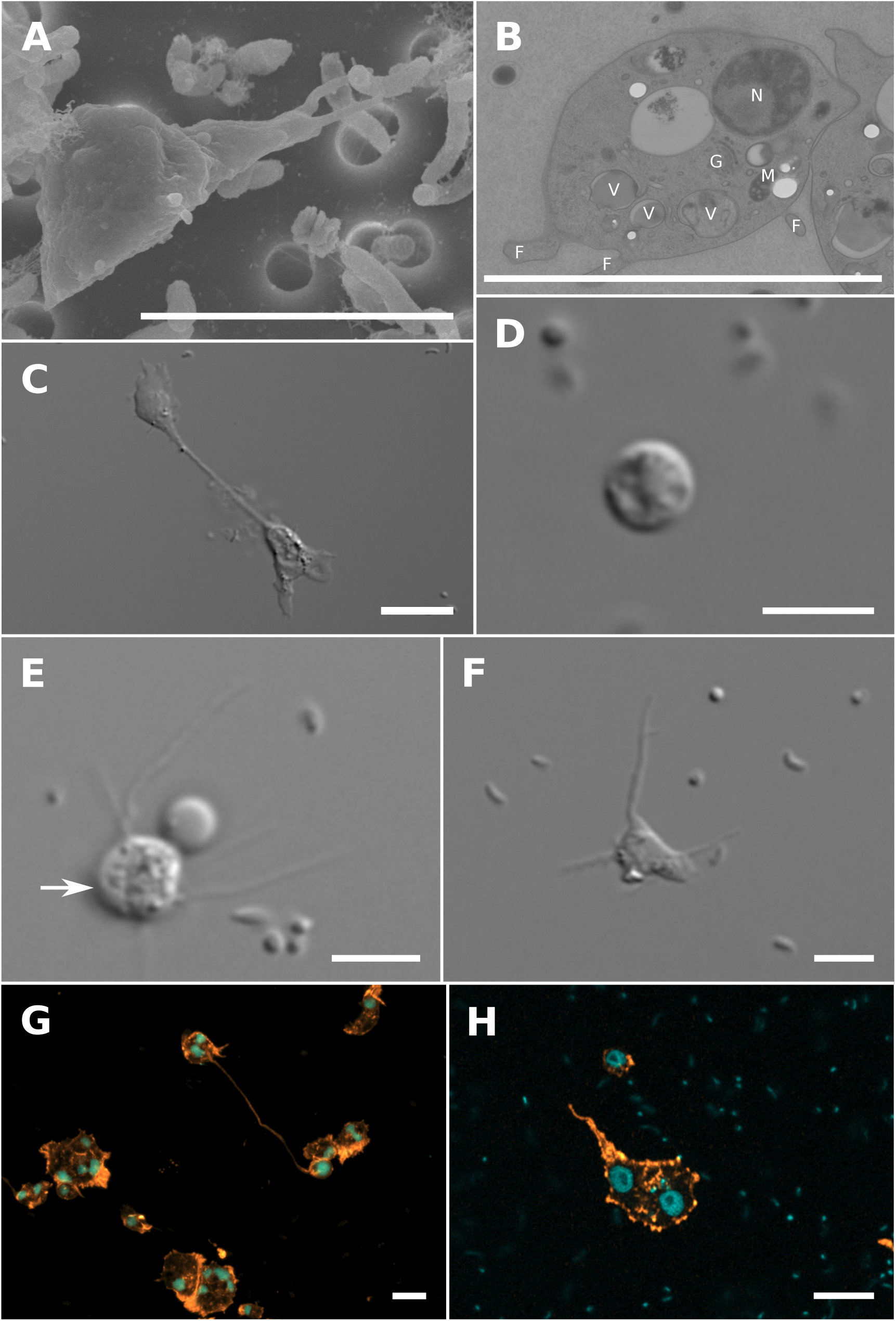
Morphology and ultrastructure of BEAP0066. A. Scanning electron microscopy of a single trophozoite cell with multiple morphologically diverse prey bacteria. B. Transmission electron microscopy of a trophozoite cell (N = Nucleus, V = Vacuole, F = Filopodia, M = Mitochondrion, G = Golgi). C. Differential interference contrast (DIC) image of the “double-amoeba” morphology (bipolarization of a cell and the generation of two poles that coexist and act semi-independently). D. DIC image of a cyst (Supplementary video 1 contains videos of the encystment process). E. DIC image of intermediate stage to encystment. Arrow points to encysting cell. F. DIC image of a trophozoite (see Supplementary video 1 for multiple instances of trophozoite behavior and movement). G and H. Immunofluorescence images stained with Phalloidin 488 (actin = orange) and Draq-5 (nuclei = cyan), showing multinucleate trophozoites and the double-amoeba morphology. Scale bars in all panels = 5 μm.

BEAP0066 is also able to generate cysts of 2 - 5 μm in diameter (average 3.3 μm, n = 20) under stress conditions, such as desiccation or depletion of nutrients (Figure 2D, Supplementary video 1). It is capable of generating an intermediate stage to encystation that differs from both the trophozoite and the cyst, which we hypothesize occurs during the switch between these two life stages. This intermediate stage presents a much rounder and less flat shape than the trophozoite, with multiple filopodia, and a cell diameter that varies between 3 - 5 μm (average 3.7 μm, n = 20) (Figure 2E, Supplementary video 2). This intermediate stage to encystation displays the possibility of cell-to-cell communication by the extension of filopodia from one cell to another (Supplementary video 3).

BEAP0066 also presents a behavior similar to the emergence of cytoplasmic arms observed in Vampyrellida (Rhizaria) (Berney et al., 2013) and the “Salvador Dalí” morphotype in Flabellula (Tubulinea, Amoebozoa) (Fenchel, 2010), among others. For convenience, in this manuscript we refer to this behavior as the “double-amoeba”. The double-amoeba consists of a bipolarization of a multinucleated cell and the generation of two poles that coexist and act as independent cells for a period of time, which is always followed by a final reabsorption (Figure 2C and supplementary video 4). Although the double-amoeba behavior could be found in various cell culture conditions, it could be reliably induced by using low-nutrient medium.

Lastly, and in addition to the already commented behaviors, BEAP0066 is able to create a social structure that we refer to in this manuscript as “colonizing rings” (Figure 3). These colonizing rings are composed of a front of amoebae advancing together with a fraction of amoebae that remain inside the generated structure, which counterintuitively are the most active ones, in terms of movement, in the structure (Supplementary video 5) (Figure 3C). The exact conditions that induce this behavior have not been found, but we observed that a low number of initial amoeba cells and a pre-established bacterial mat could increase the probability of triggering the colonizing rings. It is important to remark that when this behavior is induced, the vast majority of cells in the culture flask participate in the formation of colonizing rings, which supports the hypothesis that this behavior may be coordinated. We observed that colonizing rings have the ability to fuse with each other, resulting in a larger ring (Figure 3D). Finally, bacterial density is consistently lower inside versus outside the rings (Figure 3C).

**Figure 3.**
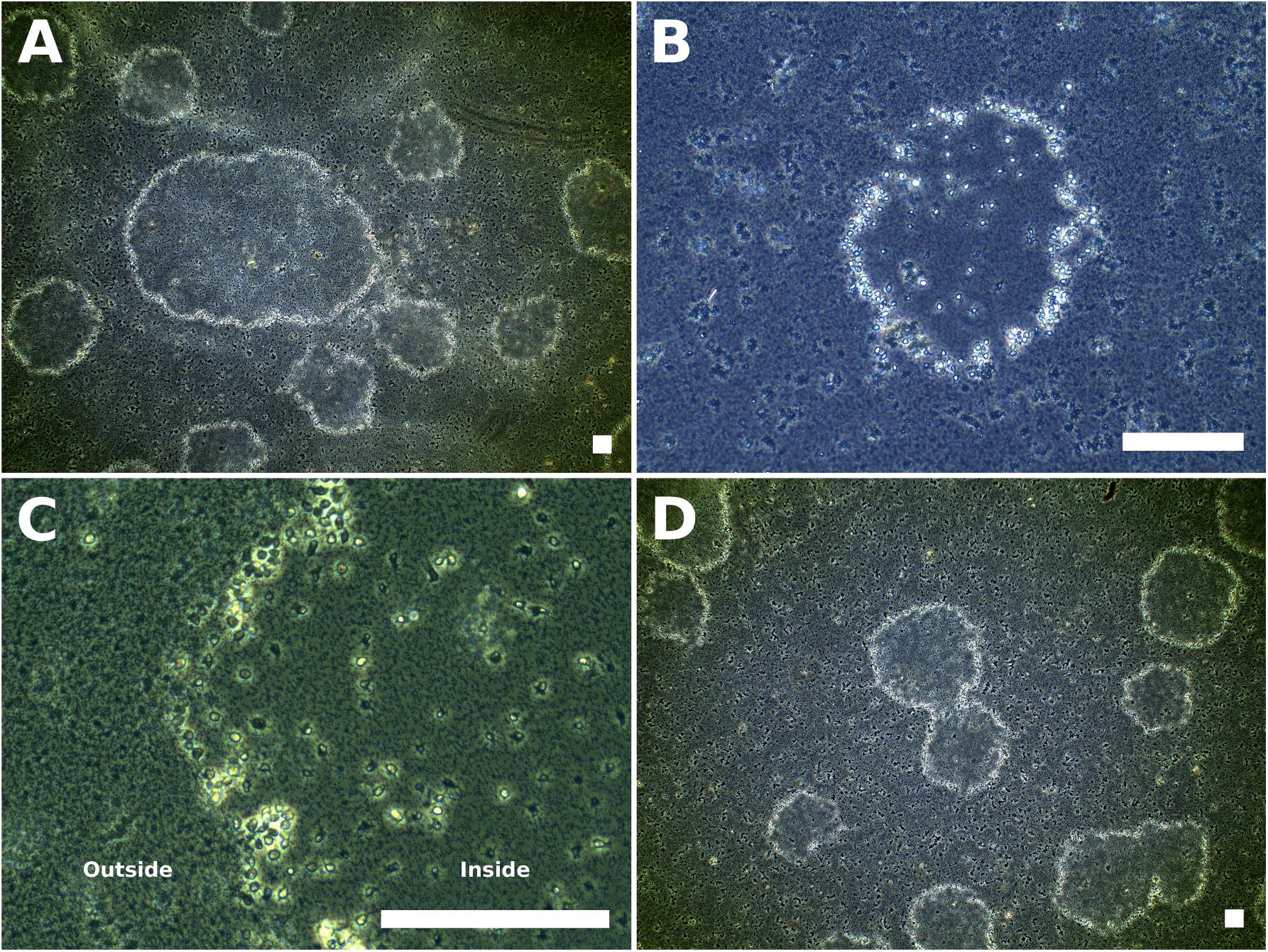
Multicellular aggregate “ring” morphology in BEAP0066. A. Multiple rings, with sizes ranging from 187 μm to 1625 μm. Also visible are numerous single amoeba cells and prey bacteria. B. Close up image of a single ring. C. Advancing front of a ring (the ring is only partially visible in this image). Bacterial density is lower inside versus outside of the ring (see Supplementary video 5). D. Two merging multicellular rings. All panels are phase contrast microscopy with scale bars = 125 μm.

### Correspondence analysis of Clusters of Orthologous Groups content in Amorphea

A recent study introduced an approach to compare evolutionary trajectories of gene content between major lineages (Ocaña-Pallarès et al., 2022). Clusters of Orthologous Groups (COG) annotation of the studied species’ proteomes allows the calculation of the relative COG content of each proteome, which is consequently used to perform a correspondence analysis with the objective of clustering species with similar relative compositions followed by projection of COG categories onto the correspondence analysis to discover associations between categories and clusters of species. COGs are functional categories of proteins grouped based on orthology that are thought to represent conserved functional roles across species. CA is a statistical method for visualizing relationships between rows and columns in a contingency table, similar to Principal Component Analysis but suited for compositional data, like the relative COG content that we analyzed. In *Ocaña-Pallarès et al*., fungal and animal species COG complement clustered independently, leading to the inference of divergent functional trajectories for Fungi and Metazoa. COG categories enriched in fungal evolution were related to metabolism, and metazoan evolution was characterized by specialization towards genes potentially related to animal multicellularity(Ros-Rocher et al., 2021; Suga et al., 2013).

In our study, we decided to extend the *Ocaña-Pallarès et al.* method to analyze functional evolution by adding Amoebozoa, a large and diverse sister group to animals and fungi. Additionally, we performed an ancestral reconstruction with orthologous groups generated with all extant species’ proteomes in order to recover the COG profile of Amoebozoa, Fungi and Metazoa common ancestors, so as to produce a more complete view of the evolution of functional content in Amorphea.

The correspondence analysis shows a clear separation of Amoebozoa, Fungi and Metazoa clusters (Figure 4A, Supplementary figure 3). This was also reflected in comparisons between each pair of groups (Supplementary figure 4). We note that 2 of the 54 amoebozoans included in the analysis, the social amoeba *Dictyostelium discoideum* and the amoebozoan *Heterostelium pallidum* are positioned more closely to Fungi than to Amoebozoa within the correspondence analysis; notably, these are two amoebozoans in our sampling with life histories that include differentiated multicellular stages. Different COG categories appear to be associated with the clusters for different lineages: The Amoebozoa cluster is associated with a higher proportion of the M and N COG categories (Cell wall/membrane/envelope biogenesis and cell motility), whereas the fungal gene repertoire was more associated with COGs with functions in metabolism (C, E, F, G, H, I and Q among others) and Metazoa presents a very strong association with the W COG category (extracellular structures; Figure 4B, Supplementary Figure 4B, Supplementary Figure 5A). Inside Amoebozoa, there is no clear separation among the Discosea, Tubulinea and Evosea lineages; their divergence might not be reflected in COG categories and therefore not detected with our methodology (Supplementary Figure 6), although we observed hints of conserved evolution of gene repertoire at lower taxonomic levels (Taxogroup2) (Berney et al., 2017) (Supplementary Figure 7).

**Figure 4.**
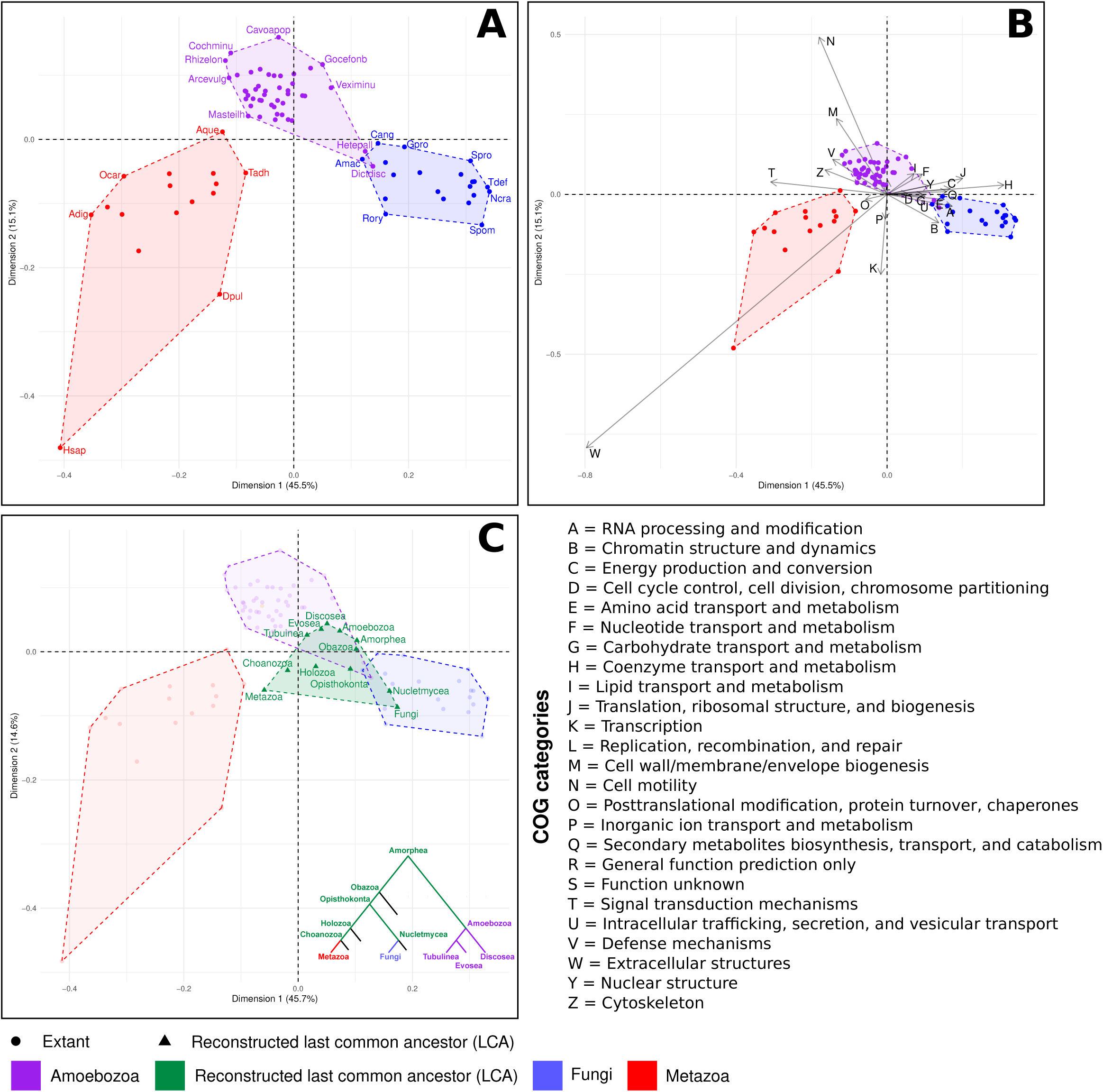
Correspondence Analyses (CA) of Clusters of Orthologous Groups (COG) functional category compositions of Amoebozoa, Metazoa, Fungi and their last common ancestors. A. First two dimensions of CA. Each point represents a single species (genome or a transcriptome; Supplementary figure 5B shows that data source does not have a significant impact). Selected points are labeled with species abbreviations: Metazoa (Adig = Acropora digitifera, Aque = Amphimedon queenslandica, Dpul = Daphnia pulex, Hsap = Homo sapiens, Ocar = Oscarella carmela and Tadh = Trichoplax adhaerens), Fungi (Amac = Allomyces macrogynus, Cang = Catenaria anguillulae, Gpro = Gonapodya prolifera, Ncra = Neurospora crassa, Rory = Rhizopus oryzae, Spom = Schizosaccharomyces pombe, Spro = Sporobolomyces roseus and Tdef = Taphrina deformans) and Amoebozoa (Arcevulg = Arcella intermedia, Cavoapop = Cavostelium apophysatum, Cochminu = Cochliopodium minutoideum, Dictdisc = Dictyostelium discoideum, Gocefonb = Gocevia fonbrunei, Hetepall = Heterostelium pallidum, Masteilh = Mastigella eilhardi, Rhizelon = Rhizomastix elongata and Veximinu = Vexillifera minutissima). Supplementary figure 3 shows the same plot with all points labeled with their corresponding species name. B. Identical CA to panel A, with individual COG category projections. C. CA with the addition of last common ancestors (green) inferred by orthologous gene identification (OrthoFinder2) and Wagner parsimony (Notung). The relationship among last common ancestors is indicated in a cartoon tree (inset).

In our ancestral reconstruction (Figure 4C), the Amorphea last common ancestor (LCA) is located between the Amoebozoa and Fungi clusters. The Amoebozoa LCA is very similar to the Amorphea LCA, retaining a higher content of M and N categories (cell membrane biogenesis and cell motility) than Metazoa and Fungi LCAs, which underwent a decrease in these categories (Supplementary Figure 8A). The Metazoa LCA falls outside the cluster of extant metazoan species, supporting a divergent genomic trajectory within metazoan evolution. Within Amoebozoa, the Tubulinea LCA appears to be more divergent from the Amoebozoa LCA than the Evosea or Discosea LCAs (Figure 4C, Supplementary Figure 8B). This analysis provides a link between COG functional evolution and phylogeny, as the reconstructed LCAs that we analyzed approach extant species as their phylogenetic position approaches the leaves of the tree.

We performed several *in silico* controls in order to validate the integrity of the results. *Ocaña-Pallarès et al.* approach used only proteomes derived from genomic data, but in order to obtain sufficient Amoebozoa sampling we also used proteomes derived from transcriptomes. Transcriptomes may be more incomplete and prone to errors than genomes; nevertheless, we found that, for the subset of species that had both genomes and transcriptomes available, the resulting clusters in the correspondence analysis were not significantly different between the two datasets, with the transcriptome gene repertoire for some species actually clustering more tightly with other species in the same lineage than the corresponding genome (Supplementary Figure 5). Next, ancestral reconstruction results could be a reflection of the inherent bias of the method used for reconstruction (Gàlvez-Morante et al., 2024). We found consistent results with two different ancestral reconstruction methods, Wagner parsimony and maximum likelihood (Supplementary Figure 9) (K. Chen et al., 2000; Guéguen et al., 2013). Lastly, we provided evidence that the clustering of Amoebozoa is driven by a biological signature and not simply by the fact that they are not Fungi or Metazoa, after introducing fake species with random COG composition that grouped completely outside of the biological clusters (Supplementary Figure 10).

### Identification of overrepresented Pfam domain clans

Our results from the correspondence analysis of Clusters of Orthologous Groups content in Amorphea shed light into the evolution of the studied supergroups. However, the limited number of COG categories makes it challenging to generate precise biological interpretations. In this context, we decided to perform a second complementary analysis based on Pfam families, as they represent structural, evolutionary and functional information. We focused on Pfam families grouped together at the clan level to further classify that information by function. Pfam clans are groups of related Pfam families (also known as Pfam domains) sharing evolutionary, structural, or functional relationships. They address limitations of sequence-based family classifications by integrating deeper evolutionary connections and structural similarities that might not be evident through sequence alone (Finn et al., 2006). At the same time, the usage of Pfam clans instead of Pfam families is a better representation of function as Pfam families often split proteins that perform the same function into separate groups, obscuring functional relationships between proteins that have evolved differently but still carry out the same role.

We performed an orthology inference for all proteins in the studied extant species with OrthoFinder2 and hierarchically clustered orthogroups by the number of representative genes present in each species. Next, we automatically defined clusters and selected groups of orthogroups specific to each lineage by cutting the dendrogram, in order to generate phylogenetically coherent orthogroup clusters. Last, we translated orthogroup presence within each detected cluster into Pfam clan content, allowing us to determine the relative abundance of Pfam clans.

In addition to a cluster of orthologous gene families present in all analyzed species, the presence/absence of orthogroups in extant species, showed clear clusters of genes specific to Amoebozoa, Fungi and Metazoa (highlighted in different colors on the vertical axis of Figure 5A). In addition to clusters of orthologs estimated to be present in phylogenetically related groups, we also observe shared clusters between taxonomic groups. There are very few genes in clusters exclusive to Fungi and either Metazoa or Amoebozoa, but there is a clear set of genes largely present in Metazoa and Amoebozoa, while absent in Fungi, which is evidence that these genes may have been lost in Fungi after their divergence from animals.

**Figure 5.**
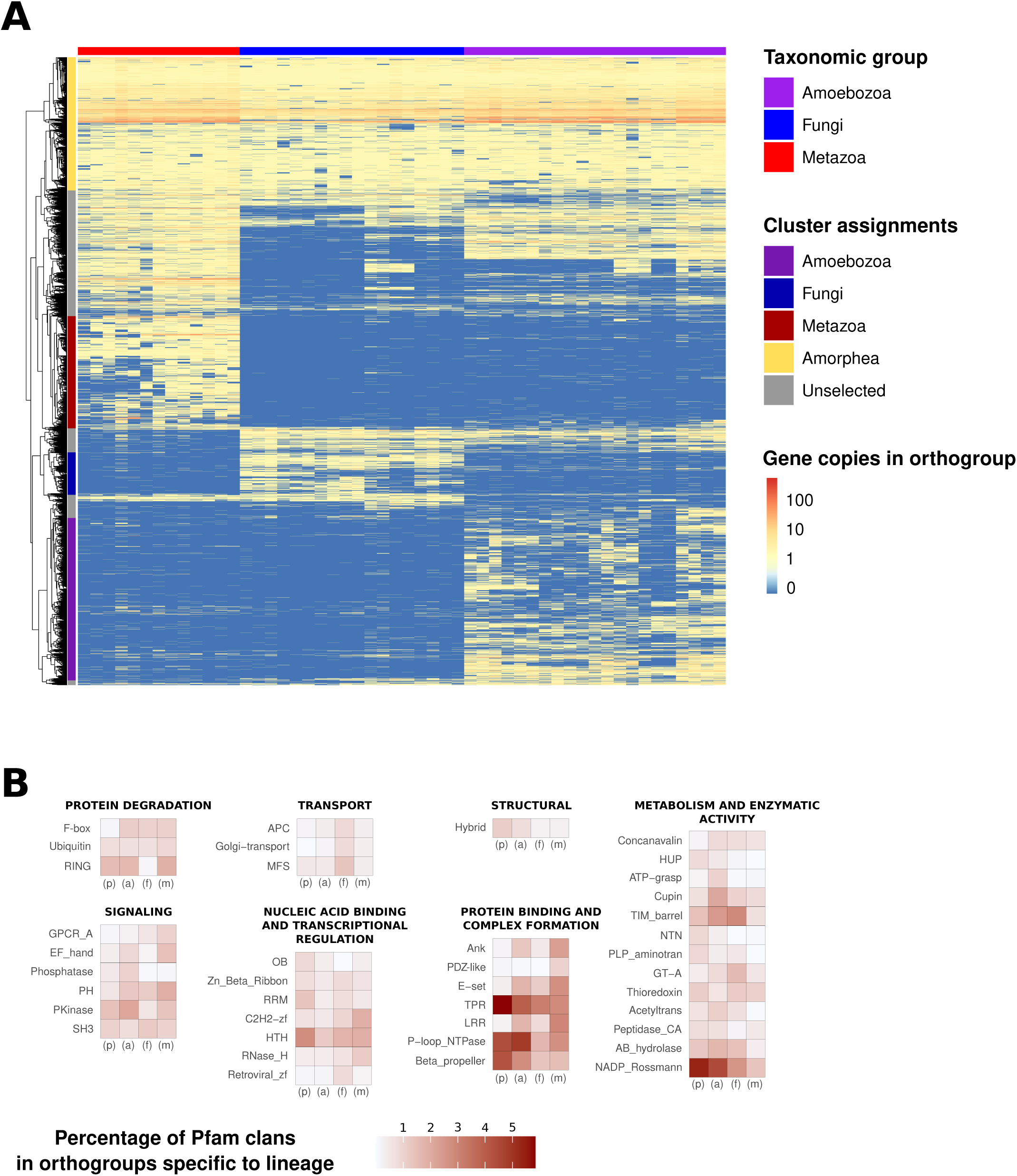
Evolution of orthologous gene families and Pfam clans in Amorphea. A. Clustering analysis (vertical axis) of gene counts for orthogroups identified in species with genomes or transcriptomes of high completeness (BUSCO eukaryota_odb10 >80%, horizontal axis). Gene counts were log10 normalized. Colored vertical bars to the left of the heatmap indicate manual assignments of hierarchical clusters as being specific to a taxonomic lineage. B. Relative proportions (expressed as percentages) of Pfam clans present in orthologous gene families assigned to specific lineages (p = Panamorphea, a= Amoebozoa, f = Fungi and m = Metazoa). Rows represent percentages of Pfam clans present in all gene families assigned to a lineage, for all Pfam clans present in the top 20 most abundant in any of the four lineages.. Pfam clans are grouped by primary/most frequent function. The Pfam domain Mito_carr is not assigned to a clan and is listed by its domain name..

To explore the functions represented in lineage-specific clusters, we first translated the presence of each orthogroup into the corresponding Pfam clans for each cluster (see Methods). Next, we combined the 20 most represented Pfam clans across all clusters to generate a heatmap of Pfam presence. To identify which Pfam clans were overrepresented in a given taxonomic group, we applied a simple heuristic: a Pfam clan was considered ‘overrepresented’ if it (1) had at least 0.5% presence in that cluster (i.e., it was not trivially present), and (2) showed a minimum 1.4-fold higher presence compared to the second-highest cluster for that same Pfam clan (i.e., it was 40% more frequent) (Figure 5B). We found the following Pfam clans to be overrepresented in the common Panamorphea cluster: Beta_propeller and TPR (whose primary functions include protein binding and complex formation); HTH, OB and RRM (Nucleic acid binding and transcriptional regulation); HUP, NTN and PLP_aminotran (Metabolism and enzymatic activity); and Hybrid (Structural). The overrepresented Pfam clans in the Amoebozoa-specific cluster were PKinase and Phosphatase (Signaling); and ATP-grasp, Acetyltrans and Cupin (Metabolism and enzymatic activity). In Fungi, GT-A (Metabolism and enzymatic activity); Retroviral-zf (Nucleic acid binding and transcriptional regulation); APC, MFS and Golgi-transport (Transport) were overrepresented. And, in Metazoa, LRR, E-set, Ank and PDZ-like (Protein binding and complex formation); EF hand, PH and GPCR_A (Signaling); and C2H2-zf and RNase_H (Nucleic acid binding and transcriptional regulation).

## DISCUSSION

BEAP0066 has a very interesting phylogenetic position and represents a new amoebozoan lineage that has been hidden in environmental samples until its characterization by light and electron microscopy, coupled with transcriptome sequencing and phylogenomics. This newly discovered lineage, with strong phylogenetic support, includes numerous environmental sequences representing diversity outside currently described amoebozoan lineages (Supplementary Figure 1). The high divergence between this new lineage and other Amoebozoa, along with the low abundance of its representatives, may suggest the existence of intermediate species that are still left to be discovered.

The intriguing behaviors of BEAP0066 that we observed could reveal new insights in Amoebozoa cell biology. We hypothesize that it could be possible that the double-amoeba behaviour results when an amoeba becomes multinucleated and the different nuclei in the cell act as local translation hubs, providing contradicting orders to the local cytoskeleton, which eventually leads to different parts of the cell moving in different directions in an uncoordinated manner. This behavior could also be a way to explore more terrain in conditions of low food availability. There are similar behaviors in Flamella (Glotova & Smirnov, 2017), which generate plasmodia that end up dividing, and Flabellula (Fenchel, 2010), which are able to create a resting stage characterized by flat multinucleated cells that can rapidly multiply in certain conditions, but the closest behavior is found in Vampyrellids (Berney et al., 2013), which extend cytoplasmatic arms to hunt for prey, and specially the behavior described for the expanded morphotype of *Leptophrys vorax* (Hess et al., 2012), which generates cells that can be drawn out to considerable length appearing as two cell bodies just connected via a thin cytoplasmic stand. In the case of the colonizing rings, we speculate that this behavior might be a strategy for the colonization of surfaces that are already covered by bacteria. We observed two behaviors of cells in rings: there are physically aggregated cells creating an advancing front of amoeba that enlarge the ring, and a second subset of cells that devour bacteria that become trapped inside the ring, generating a differential observed bacterial density when comparing the inside and outside of the rings. When it comes to the possibility of cell-to-cell communication, even though there is no further data and the following is speculation, the observed structure is superficially similar to tunneling nanotubes, which are cell bridges that mediate the intercellular transfer of organelles, plasma membrane components and cytoplasmic molecules, which have been studied mainly in Metazoa (Gerdes & Carvalho, 2008).

In our analysis of gene repertoire composition, Amoebozoa, Metazoa and Fungi showed three distinct evolutionary trajectories. Initially, we calculated the relative composition of COG categories in proteomes of extant species in Amoebozoa, Metazoa and Fungi. Next, we used ancestral reconstruction to estimate the COG profiles of their Amorphea ancestors. These data enabled us to perform correspondence analysis to explore differences in COG composition based on the taxonomy of extant and reconstructed species. Our findings align with previous studies (Ocaña-Pallarès et al., 2022), replicating associations with COGs related to animal multicellularity in the case of Metazoa and COGs related to metabolism in the case of Fungi. In contrast, the addition of new data in our study reveals that Amoebozoa display a third evolutionary trajectory enriched in the M (cell wall/membrane biogenesis) and N (cell motility) COG categories, likely to be related to the amoeboid morphologic flexibility, crucial for motility, phagocytosis and generation of various life stages and structures. The reconstruction of COG profiles of Amorphea ancestors revealed intriguing insights into evolutionary relationships. Our findings suggest that the ancestral Amoebozoa proteome is closer than the last common ancestor of Metazoa or Fungi to the last common ancestor of Amorphea, with relatively few changes in a period of evolutionary stasis, in terms of the distribution of genomic functions among COG categories. In contrast, metazoan extant species exhibit a high degree of divergence from the Metazoa last common ancestor, reflecting a continued evolution during the birth of the different lineages within Metazoa.

To add to our conclusions based on Amorphea relative COG content, we decided to use Pfam family clans in a complementary analysis. COG categories are a useful tool that give us a general overview on functional content, but Pfam clans offer more specific functional detail than COGs. Pfam clans are more suited to our study than plain Pfam families (also known as domains), as Pfam clans group together families of proteins that perform the same function. We gathered all proteins in the studied extant species, inferred orthologous gene families and hierarchically clustered them. We detected putative Amoebozoa-specific, Fungi-specific and Metazoa-specific clusters, as well as a cluster of orthologous gene families present in all extant species analyzed, and several clusters with more complex evolutionary histories. Next, we translated protein presence within each cluster of orthologous gene families into Pfam clan content, allowing us to determine the relative abundance of Pfam clans. In the case of Metazoa we can again detect a specialization towards animal multicellularity with Pfam clans like LRR or E-set, which are often found in genes with functions in cell-cell adhesion and the animal immune system, and PDZ-like, which is critical for the assembly of supramolecular complexes and signal transduction (Jeleń et al., 2003; Sheng & Sala, 2001). Fungi showed a higher specialization towards Pfam clans related to transport, such as MFS, APC and Golgi-transport, which could be essential for the uptake and secretion of enzymes, nutrients and metabolites of the fungal lifestyle. The detection of a large shared cluster between Amoebozoa and Metazoa, together with the fact that the Fungi specific cluster is clearly smaller than the others, are consistent with observations that gene loss played a role in fungal evolution (Merényi et al., 2023). In Amoebozoa, we could detect a specialization in phosphate-mediated signal transduction, with a diversification of protein kinases and phosphatases, which together with the high content in ATP-grasp could be related to the need for a rapid response to diverse environmental stimuli.

This project identified distinct evolutionary trajectories within Fungi, Metazoa, and Amoebozoa by analyzing patterns in COG categories and Pfam family clans. These findings support the hypothesis that the phylogenetic relationships among the lineages are reflected in the relative compositions of their gene repertoires. We hypothesize that the evolutionary trajectory of Amoebozoa, shaped by the detected COG categories and Pfam clans, retained a partially ancestral eukaryotic state optimized for their particular predatory lifestyle. This state was centered on phagocytosis, motility, and cellular plasticity, enabling rapid responses to environmental signals. This could have enabled access to bacterial mats, the primary food source in the mid-Proterozoic biosphere, the era in which Amoebozoa first emerged (Tekle et al., 2022), and made Amoebozoa highly adaptable to fluctuating conditions, reinforcing their ecological success.

For future studies, we propose some follow up experiments to test our hypotheses. A transcriptomic analysis of Amoebozoa extant species under different conditions favouring or limiting phagocytosis (presence/absence of prey, low/high temperature, viscous/liquid medium…) could be performed in order to detect genes specifically involved in amoebozoan predation. After the identification of those genes, we could make use of the data generated in this project to measure the relative proportions of these genes in Fungi, Metazoa, Amoebozoa and the reconstructed Amorphea ancestor, to confirm that the genes that Amoebozoa retained from the last common ancestor of Amorphea are indeed differentially regulated in predation. We also propose to test Amoebozoa environmental adaptability by exposing cultures to different laboratory conditions and to monitor changes in morphology and gene expression, as well as the limits of adaptation to each certain studied variable (temperature, pressure, nutrient availability, inhibition vs. stimulation of attachment to surfaces). As we hypothesized that Amoebozoa developed a evolutionary strategy based on rapid environmental response; due to its content in kinases, phosphatases and ATP-grasp; we could check if there is an upregulation of such proteins in “stress” conditions, in order to elucidate if they confer an advantage in terms of environmental adaptability. All described potential experiments should be performed with appropriately matched non-amoebozoan control species that would let us extract Amoebozoa-specific conclusions.

We believe that testing our hypothesis will enable a more comprehensive characterization of these evolutionary trajectories. By studying and identifying a broader range of functional evolutionary trajectories, we aimed to deepen our understanding of the molecular and functional innovations underlying eukaryotic cell evolution. This knowledge will provide critical insights into the diversity and complexity of life at the cellular level.

## TAXONOMIC SUMMARY

Eukaryota, Amorphea, Amoebozoa, Discosea, Centramoebia

Apostamoebida Gàlvez-Morante and Richter, n. ord.

Description: The most inclusive clade containing the genus *Apostamoeba* but excluding the clades Pellitida, Himatismenida and Acanthopodida.

Etymology: Named after the Greek term “Apostolos”, meaning “the one who is sent away” or “messenger”, in reference to its high phylogenetic divergence.

Zoobank registration: Described under the Zoological Code; Zoobank registration will be performed following peer review.

Apostameobidae Gàlvez-Morante and Richter, n. fam.

Description: Apostamoebida species, characterized by an 18S rRNA genetic signature matching the following: sequence “TCACTTTGAATAAATTG” at position 759 - 775 of *Apostamoeba explorator* 18S (localized in the region helix E23_8), and consensus sequence “CATTTTAGCATAGGATGA” at position 811 - 828 of *Apostamoeba explorator* 18S with one allowed discrepancy (localized in the region helix E23_9) (Wuyts et al., 2000).

Etymology: Named after the Greek term “Apostolos”, meaning “the one who is sent away” or “messenger”, in reference to its high phylogenetic divergence.

Zoobank registration: Described under the Zoological Code; Zoobank registration will be performed following peer review.

*Apostamoeba* Gàlvez-Morante and Richter, n. gen.

Description: Trophozoites range from 4-17 μm long (average 7.9 μm) and 2-7 μm wide (average 2.3 μm), producing a length-to-breadth ratio of 1-5 (average 2.3), while their nuclei measure 1-2.5 μm (average 1.7 μm); their size varies with growth conditions like temperature, nutrient levels, and prey density. The locomotive stage is flat with multiple, variably sized filopodia that can extend several cell lengths, and larger cells are often multinucleated. Under stress, they form cysts (2-5 μm, average 3.3 μm) and may enter an intermediate, rounder encystation stage (3-5 μm, average 3.7 μm).

Etymology: Named after the Greek term “Apostolos”, meaning “the one who is sent away” or “messenger”, in reference to its high phylogenetic divergence and the fact that it is the first described genus of an undiscovered vast amount of environmental diversity inside Apostamoebida”. *Apostamoeba* is feminine.

Type species: *Apostamoeba explorator* (see below).

Zoobank registration: Described under the Zoological Code; Zoobank registration will be performed following peer review.

*Apostamoeba explorator* Gàlvez-Morante and Richter, n. sp.

Description: With the characters of the genus. Additionally, they exhibit “double-amoeba” behavior, where a multinucleated cell temporarily splits into two independent poles before reuniting, and form “colonizing rings” in which groups of amoebae advance together with some remaining behind.

Etymology: The latin term “explorator” refers to its ability and behaviors to investigate its surroundings.

Type locality: Subsurface water, Mediterranean Sea, Blanes Bay, Spain, 41°40’ N / 2°40’ E.

Type material: The name-bearing type (an hapantotype) is an SEM stub deposited in the Marine Biological Reference Collections (CBMR) at the Institut de Ciències del Mar (ICM-CSIC, Barcelona, Spain) under the catalog/accession number ICMCBMR000695 (Guerrero et al., 2023). This material also contains uncharacterized prokaryote species, which do not form part of the hapantotype.

Gene sequence: The rRNA gene sequence (18S, ITS1, 5.8S, ITS2, 28S) of strain BEAP0066 is deposited in Genbank as PV590302.

Cell culture: A culture containing BEAP0066 and mixed bacterial species is publicly available and has been deposited in the Roscoff Culture Collection (March, 2025; currently awaiting assignment of RCC ID).

Zoobank registration: Described under the Zoological Code; Zoobank registration will be performed following peer review.

This publication (work) will be registered with Zoobank following peer review.

## METHODS

### Isolation and culture

BEAP0066 was isolated from the coast of Spain, in Blanes Bay (41°40’ N / 2°40’ E) in November 2021 through the dilution of surface marine water in RS medium (1 NM : 10 NNM concentration). RS medium is a fully defined medium designed to grow a wide range of marine protists. It is composed of two components: the nutrient medium (NM) and the non-nutrient medium (NNM) (https://mediadive.dsmz.de/medium/P4, https://mediadive.dsmz.de/medium/P5) (Sigona *et al*., in preparation).

BEAP0066 was maintained in 25 cm^2^ flasks with a total volume of 10 mL, with a split frequency of 1 split per 1.5 weeks at 18 °C, in the presence of light. The splits were performed by adding 9 mL of R/S medium (1 NM : 10 NNM concentration, also viable at 1:50 and 1:100) and 1 mL of the previously growing culture, with scraping.

*Vannella septentrionalis* BEAP0079 was isolated from the coast of Spain, in Blanes Bay (41°40’ N / 2°40’ E) in 2021 through the dilution of surface marine water in RS medium (1 NM : 10 NNM concentration).*V. septentrionalis* was maintained in 25 cm^2^ flasks with a total volume of 10 mL, with a split frequency of 1 split per 1.5 weeks at 12 °C, in the absence of light. The splits were performed by adding 9 mL of R/S medium (1 NM : 50 NNM concentration, also viable at 1:10 and 1:100) and 1 mL of the previously growing culture, with scraping.

### 18S rRNA gene sequencing

A sample of 50 mL (two 75 cm^2^ flasks with a total volume of 25 mL) was used for 18S cloning. The cells were centrifuged for 20 minutes at 13000 x g and 4 °C. DNA was extracted using the DNeasy PowerSoil Pro kit (QIAGEN) and the 18S rRNA gene was amplified by gradient PCR (from 47 °C to 58 °C) using a vannelid-specific forward primer together with a universal eukaryotic reverse primer (CCATGCAAGTCTAAGTATAAATCAT - CCTTCYGCAGGTTCACCTAC) (López-García et al., 2003; Vaulot et al., 2022). Amplification products were purified using NZYGelpure kit (NZYTech), cloned using the TOPO-TA Cloning kit (Invitrogen) and transformed into *E. coli* cells following LacZα-complementation. Positive clones were selected and amplified by PCR with vector-specific primers M13F-M13R (GTAAAACGACGGCCAGT - CAGGAAACAGCTATGAC). Sequencing was performed by Eurofins genomics using Sanger sequencing. The resulting sequences were base called using phred (Ewing et al., 1998) with the parameters ‘-trim_alt “” -trim_cutoff 0.01’ and assembled with phrap (de la Bastide & McCombie, 2007) with the parameter ‘-repeat_stringency 0.4’ and the consensus sequence was exported with consed (Gordon & Green, 2013). Assembled sequences were identified through BLAST versus EukRibo (Berney et al., 2022).

While the 18S sequences obtained with these methodologies were used to initially identify the two species, for 18S phylogenetic trees and upload to NCBI we instead used the sequences assembled by phyloFlash (Gruber-Vodicka et al., 2020) for BEAP0079 and Trinity (Grabherr et al., 2011) for BEAP0066 (contig TRINITY_DN167_c0_g1_i10). See the process below.

### RNA extraction

In order to extract and sequence total RNA we used a similar experimental approach as the one described by Richter and colleagues (Richter et al., 2018). Prior to RNA isolation, both strains were grown in large batches using two 75 cm2 Rectangular Canted Neck Cell Culture Flasks with Vented Caps (353136, Corning Life Sciences), each containing 50 ml of medium.

We isolated total RNA from both strains by using the RNAqueous kit (AM1912, ThermoFisher Scientific). RNA concentration was measured using a NanoDrop One/One C UV-Vis Microvolume Spectrophotometer (ThermoFisher Scientific). We digested genomic DNA using the TURBO DNA-free kit (AM1907, ThermoFisher Scientific) according to the manufacturer’s instructions and removed DNase using DNase Inactivation Reagent. The quality evaluation of extracted total RNA, the library preparation, and the sequencing were carried out at the CRG Genomics Core Facility in Barcelona. The quality of extracted total RNA was assessed by Bioanalyzer 2100 RNA Pico chips (Agilent Technologies). To prepare stranded libraries, the NEBNext Ultra II Directional RNA Library Prep Kit Total RNA was used, applying double the normal rounds of PolyA selection prior to cDNA synthesis to reduce the excess of bacterial RNA. Finally, the RNA was sequenced by using Illumina NextSeq 2000 (2x150 bp).

### *De novo* transcriptome assembly and proteome generation

The quality assessment of the sequenced RNA was performed using FastQC v0.11.9 (Andrews S., 2010), while evaluating the purity of the sequences using phyloFlash v3.4 (Gruber-Vodicka et al., 2020), a tool designed to assess the 16S/18S rRNA gene taxonomic composition of metagenomic datasets. The elimination of artifacts and adapter sequences was achieved via fastp v0.23.2 (S. Chen et al., 2018) with the following parameters:

--low_complexity_filter, --cut_front, --cut_tail, --cut_right, --cut_front_window_size 1

--cut_tail_window_size 1, --cut_right_window_size 4, --cut_mean_quality 5, --trim_front1 12

--trim_front2 12, --trim_poly_g, --trim_poly_x

--adapter_sequence=AGATCGGAAGAGCACACGTCTGAACTCCAGTCA

--adapter_sequence_r2=AGATCGGAAGAGCGTCGTGTAGGGAAAGAGTGT. *De novo* transcriptome assembly was performed using Trinity v2.14.0 (Grabherr et al., 2011) with parameter --SS_lib_type RF.

We decontaminated the newly assembled transcriptome by blastn against a set of potential contaminant genomes (species with a 16S/18S detected by phyloFlash or Trinity; Supplementary file 3) and removing matching contigs with a percentage identity of ≥ 98% and match length > 100. WinstonCleaner with default parameters was also used to remove cross-contamination between samples sequenced in the same run (https://github.com/kolecko007/WinstonCleaner). The FCS-adaptor v.0.5.5 tool was also run to further decontaminate our transcriptomes, which cleaned 2 contigs from BEAP0066 and 3 contigs from BEAP0079 (Astashyn et al., 2024).

Subsequently, protein prediction was performed using TransDecoder v5.5.0 with the ‘-S’ option for strand specificity (https://github.com/TransDecoder/TransDecoder) and Pfam protein annotation was carried out with InterProScan 5.56-89.0 with default parameters except for --disable-precalc and --applications Pfam, Phobius (Blum et al., 2021). To examine the completeness of our transcriptomes, we searched for the presence of a set of conserved, single-copy orthologous genes by employing BUSCO v5.3.2 with the Eukaryota odb10 dataset (Simão et al., 2015).

### Phylogenetic tree inference

The predicted proteome of BEAP0066, together with an increased sampling for Amoebozoa (Supplementary file 2), underwent PhyloFisher’s protocol for phylogenomic assembly (Tice et al., 2021), following the standard specified instructions. Next, a maximum likelihood tree was generated with IQ-TREE, which automatically selected the LG+R10 model, with 1000 ultrafast bootstraps (Nguyen et al., 2015) (Supplementary file 1).

### Light Microscopy

For Differential Interference Contrast (DIC) microscopy, cells were seeded on a glass-bottom dish. Imaging was conducted using a Zeiss Axiovert Inverted microscope equipped with a 63x oil-immersion lens. Digital images were processed using Fiji software (Schindelin et al., 2012). For Phase Contrast microscopy cells were seeded in 6-well plates. Imaging was conducted using a Zeiss Axiovert Inverted microscope and pictures were taken with a mobile phone.

### Scanning Electron Microscopy

Cells were fixed with 2.5% glutaraldehyde for 3 hours at room temperature and then seeded on membranes with 0.8 μm pores (WHA10417301, MERCK Chemicals and Life Science) via filtering. The samples were dehydrated through a graded ethanol series and dried by critical point with liquid carbon dioxide in a Leica EM CPD300 unit (Leica Microsystems, Austria). The dried filters were mounted on stubs with colloidal silver and then were sputter-coated with gold in a Q150R S (Quorum Technologies, Ltd.) and observed with a Hitachi SU8600 field emission scanning electron microscope (Hitachi High Technologies Co., Ltd., Japan) in the Electron Microscopy Service of the Institute of Marine Science (ICM-CSIC), Barcelona.

### Transmission Electron Microscopy

Sample preparation was performed in the Electron Cryomicroscopy Unit at the Scientific and Technical Centers of the University of Barcelona. Samples were concentrated by centrifugation (2000 x g, 5 minutes) and resuspended in 20 % BSA in artificial sea water. Most of the supernatant was removed and the concentrated cells cryo-immobilized using a Leica HPM100 high-pressure freezer (Leica Microsystems, Vienna, Austria). Samples were freeze-substituted in pure acetone containing 2 % (w/v) osmium tetroxide and 0.1 % (w/v) uranyl acetate at -90 °C for 72 hours in an EM AFS2 (Leica Microsystems, Vienna, Austria). Later, they were warmed up to 4 °C at a 5 °C/h slope, kept at 4°C for 2 hours, and transferred to room temperature and kept for 2 hours in darkness. Samples were washed in acetone at room temperature, infiltrated in increasing concentrations of Epon-812 resin in acetone until purely Epon-812. Then, they were embedded and polymerized in Epon-812 at 60 °C for 48 hours. Ultrathin sections of 60 nm were obtained with a UC6 ultramicrotome (Leica Microsystems, Vienna, Austria) and placed on Formvar-coated copper grids. Sample sections were stained with 2 % (w/v) uranyl acetate for 30 minutes, lead citrate for 5 minutes and examined in a TEM Jeol JEM 1010 (Gatan, Japan) equipped with a tungsten cathode. Images were acquired at 80 kV with a 1k x 1k CCD Megaview III camera.

### Immunofluorescence microscopy

Cells were seeded on coverslips pre-treated with poly-L-lysine to enhance cell adherence to the glass surface, washed once with artificial sea water (ASW), and fixed with a solution of 4% formaldehyde in ASW for 5 minutes. Cells were washed for 5 minutes with ASW, blocked and permeabilized in a solution containing 1% BSA and 0.3% Triton X-100 in ASW for 1 hour. After, cells were incubated with 132 nM phalloidin 488 (A12379, ThermoFisher Scientific) in ASW at room temperature for 30 minutes, in order to stain actin. DNA was stained with 0.2 μM Draq-5 (62251, ThermoFisher Scientific) for 15 min. Coverslips were mounted in ProLong Gold antifade reagent (P36934, Thermo Fisher Scientific).

The observation of the samples was carried out at the Advanced Light Microscopy Unit (ALMU) located at the Centre for Genomic Regulation (CRG) in Barcelona, using a Zeiss LSM 980 confocal microscope equipped with Airyscan 2 super-resolution (405, 488, 561, 639 nm lasers) and a 63x lens. Digital images were later processed using Fiji software (Schindelin et al., 2012).

### Orthogroup generation, functional annotation and clustering

The studied proteomes were filtered with CD-HIT to remove redundancy (Fu et al., 2012) and screened for bacterial contamination using BLAST against eukaryotic and bacterial databases (Altschul et al., 1990). We used OrthoFinder2 with default parameter values, in order to cluster them into orthogroups, stored in file “Orthogroups.tsv” (Emms & Kelly, 2019).

We used eggNOG-mapper v2 with default parameter values to functionally annotate the proteomes of this project (Cantalapiedra et al., 2021). To infer the functional composition of each orthogroup, we collected all associated protein annotations and normalized their contributions by dividing each annotation by the total number of annotations within the orthogroup. If a protein had multiple functional annotations, its contribution was distributed equally among them by assigning a weight of 1 divided by the number of annotations.

The clustering of proteomes was performed in RStudio (RStudio Team, 2020) with package pheatmap (Raivo Kolde, 2010).

*Rhizophagus irregularis* was excluded from this analysis due to its artifactual behavior (Supplementary Figure 5), clustering closer to Metazoa than Fungi; which is not observed in Ocaña-Pallarès et al. although the same initial data and methodology was applied. *Rhizophagus irregularis’*s transcriptome did cluster with Fungi, which strengthened our suspicions on *Rhizophagus irregularis’*s genome. We hypothesize there was a mislabeling with *Rhizophagus irregularis’*s sequence, or that our newer version of eggNOG-mapper v2 gave us a different output (Cantalapiedra et al., 2021). Only Amoebozoa species with a high BUSCO value (> 80%) were used in the clustering.

### Correspondence analyses

Correspondence analyses were done in RStudio (RStudio Team, 2020) with the FactoMineR (Lê et al., 2008) package and the plots were constructed with the ggplot2 (Wickham, 2011) package.

*Rhizophagus irregularis* was excluded from this analysis as described above. Several Amoebozoa species (*Micriamoeba sp. ATCC-PRA-134*, *Squamamoeba japonica*, *Ovalopodium desertum*, *Mayorella cantabrigiensis*, *Thecamoeba sp. SK13-4B*, *Echinostelium bisporum*, *Mycamoeba gemmipara*, *Tychosporium acutostipes* and *Pellita catalonica*) were also excluded due to a low BUSCO value (< 40%).

### Ancestral reconstruction

A proteomic dataset of 117 species representing Amorphea diversity and outgroups was used for ancestral reconstruction (36 of these species covering Fungi and Metazoa were obtained from Ocaña-Pallarès et al.) (Supplementary file 2). Among these species we include our BEAP0066 and *Vannela septentrionalis* assembled transcriptomes.

The input tree for the ancestral reconstruction was a manual merging of PhyloFisher and IQ-TREE’s output (described above) and the Fungi and Metazoa topology from Ocaña-Pallarès et al. This topology was treated with Bppml (Guéguen et al., 2013), in order to estimate branch lengths and model parameters, with a configuration file, specifying a stationary process with a binary model and a gamma distribution of rate variation among sites, which was stated to remain homogeneous across all branches of the topology.

We ran our ancestral reconstructions with two different programs, in order to minimize and detect biased results generated by a strong effect of the methodology instead of the data (Gàlvez-Morante et al., 2024).

We ran Bppancestor (maximum likelihood, counts presence/absence of genes per orthogroup) with a configuration file specifying the most likely model estimated by Bppml. The used version was Bio++ version 3.0.0 (Guéguen et al., 2013).

We ran Notung (Wagner parsimony, counts number of ancestral genes per orthogroup) with the following command “notung -b batch_notung.txt --reconcile --events --parsable

--absfilenames --log --silent --progressbar --speciestag prefix --phylogenomics”. The used version was Notung version 2.9.1 (K. Chen et al., 2000).

*Rhizophagus irregularis* was excluded from this analysis as described above.

## DATA, CODE AND MATERIALS AVAILABILITY

Data: Transcriptome reads and assemblies are available in GenBank under BioProject accession number PRJNA1257292. The ribosomal operon sequence (18S, ITS1, 5.8S, ITS2, 28S) sequence of BEAP0066 is available with accession number PV590302, and the 18S sequence of BEAP0079 is available with accession number PV584300. Supplementary files, videos, transcriptome assemblies, predicted proteomes and input data are available in FigShare with the DOI 10.6084/m9.figshare.28741229. The transcriptome assemblies available on GenBank differ from those on FigShare due to the automatic decontamination tools used by GenBank, which we believe have erroneously removed a number of valid contigs. As a result, we recommend using our FigShare assemblies, which retain these sequences, over the GenBank versions.

Code: Scripts used in data analysis are available on GitHub at: https://github.com/beaplab/amorphea_evolution

Cell cultures: Cell cultures containing BEAP0066 (and mixed bacteria) and BEAP0079 (and mixed bacteria) are publicly available and have been deposited in the Roscoff Culture Collection (March, 2025; currently awaiting assignment of RCC IDs).

## ACKNOWLEDGEMENTS

We thank Clara Cardel’s, operating the Blanes Bay Microbial Observatory (BBMO), and Drs. Josep M. Gasol and Ramon Massana, for providing access to monthly water samples from Blanes Bay. We also thank Adrià Auladell and Eduard Ocaña-Pallarès for their intellectual contributions to the project and Matt Brown for critical comments on the manuscript. Last, we thank Tanasije Rakić for isolating *V. septentrionalis* BEAP0079.

## REFERENCES

1. Altschul, S. F., Gish, W., Miller, W., Myers, E. W., & Lipman, D. J. (1990). Basic local alignment search tool. Journal of Molecular Biology, 215(3), 403–410. 10.1016/S0022-2836(05)80360-2

2. Andrews S. (2010). FastQC: A Quality Control Tool for High Throughput Sequence Data [Online]. [Computer software]. Available online at: http://www.bioinformatics.babraham.ac.uk/projects/fastqc/

3. Astashyn, A., Tvedte, E. S., Sweeney, D., Sapojnikov, V., Bouk, N., Joukov, V., Mozes, E., Strope, P. K., Sylla, P. M., Wagner, L., Bidwell, S. L., Brown, L. C., Clark, K., Davis, E. W., Smith-White, B., Hlavina, W., Pruitt, K. D., Schneider, V. A., & Murphy, T. D. (2024). Rapid and sensitive detection of genome contamination at scale with FCS-GX. Genome Biology, 25(1), 60. 10.1186/s13059-024-03198-7

4. Berney, C., Ciuprina, A., Bender, S., Brodie, J., Edgcomb, V., Kim, E., Rajan, J., Parfrey, L. W., Adl, S., Audic, S., Bass, D., Caron, D. A., Cochrane, G., Czech, L., Dunthorn, M., Geisen, S., Glöckner, F. O., Mahé, F., Quast, C., … de Vargas, C. (2017). UniEuk: Time to Speak a Common Language in Protistology! Journal of Eukaryotic Microbiology, 64(3), 407–411. 10.1111/jeu.12414

5. Berney, C., Henry, N., Mahé, F., Richter, D. J., & Vargas, C. de. (2022). *EukRibo: A manually curated eukaryotic 18S rDNA reference database to facilitate identification of new diversity* (p. 2022.11.03.515105). bioRxiv. 10.1101/2022.11.03.515105

6. Berney, C., Romac, S., Mahé, F., Santini, S., Siano, R., & Bass, D. (2013). Vampires in the oceans: Predatory cercozoan amoebae in marine habitats. The ISME Journal, 7(12), 2387–2399. 10.1038/ismej.2013.116

7. Blum, M., Chang, H.-Y., Chuguransky, S., Grego, T., Kandasaamy, S., Mitchell, A., Nuka, G., Paysan-Lafosse, T., Qureshi, M., Raj, S., Richardson, L., Salazar, G. A., Williams, L., Bork, P., Bridge, A., Gough, J., Haft, D. H., Letunic, I., Marchler-Bauer, A., … Finn, R. D. (2021). The InterPro protein families and domains database: 20 years on. Nucleic Acids Research, 49(D1), D344–D354. 10.1093/nar/gkaa977

8. Borkens, Y. (2024). The Pathology of the Brain Eating Amoeba Naegleria fowleri. Indian Journal of Microbiology. 10.1007/s12088-024-01218-5

9. Brunet, T., Albert, M., Roman, W., Coyle, M. C., Spitzer, D. C., & King, N. (2021). A flagellate-to-amoeboid switch in the closest living relatives of animals. eLife, 10, e61037. 10.7554/eLife.61037

10. Cantalapiedra, C. P., Hernández-Plaza, A., Letunic, I., Bork, P., & Huerta-Cepas, J. (2021). eggNOG-mapper v2: Functional Annotation, Orthology Assignments, and Domain Prediction at the Metagenomic Scale. Molecular Biology and Evolution, 38(12), 5825–5829. 10.1093/molbev/msab293

11. Chen, K., Durand, D., & Farach-Colton, M. (2000). NOTUNG: A Program for Dating Gene Duplications and Optimizing Gene Family Trees. Journal of Computational Biology, 7(3–4), 429–447. 10.1089/106652700750050871

12. Chen, S., Zhou, Y., Chen, Y., & Gu, J. (2018). fastp: An ultra-fast all-in-one FASTQ preprocessor. Bioinformatics, 34(17), i884–i890. 10.1093/bioinformatics/bty560

13. de la Bastide, M., & McCombie, W. R. (2007). Assembling genomic DNA sequences with PHRAP. *Current Protocols in Bioinformatics*, Chapter 11, Unit11.4. 10.1002/0471250953.bi1104s17

14. Emms, D. M., & Kelly, S. (2019). OrthoFinder: Phylogenetic orthology inference for comparative genomics. Genome Biology, 20(1), 238. 10.1186/s13059-019-1832-y

15. Ewing, B., Hillier, L., Wendl, M. C., & Green, P. (1998). Base-calling of automated sequencer traces using phred. I. Accuracy assessment. Genome Research, 8(3), 175–185. 10.1101/gr.8.3.175

16. Fenchel, T. (2010). The Life History of *Flabellula baltica* Smirnov (Gymnamoebae, Rhizopoda): Adaptations to a Spatially and Temporally Heterogeneous Environment. Protist, 161(2), 279–287. 10.1016/j.protis.2009.10.005

17. Finn, R. D., Mistry, J., Schuster-Böckler, B., Griffiths-Jones, S., Hollich, V., Lassmann, T., Moxon, S., Marshall, M., Khanna, A., Durbin, R., Eddy, S. R., Sonnhammer, E. L. L., & Bateman, A. (2006). Pfam: Clans, web tools and services. Nucleic Acids Research, 34(Database issue), D247–D251. 10.1093/nar/gkj149

18. Fu, L., Niu, B., Zhu, Z., Wu, S., & Li, W. (2012). CD-HIT: Accelerated for clustering the next-generation sequencing data. Bioinformatics, 28(23), 3150–3152. 10.1093/bioinformatics/bts565

19. Gabr, A., Zournas, A., Stephens, T. G., Dismukes, G. C., & Bhattacharya, D. (2022). Evidence for a robust photosystem II in the photosynthetic amoeba Paulinella. New Phytologist, 234(3), 934–945. 10.1111/nph.18052

20. Gàlvez-Morante, A., Guéguen, L., Natsidis, P., Telford, M. J., & Richter, D. J. (2024). Dollo Parsimony Overestimates Ancestral Gene Content Reconstructions. Genome Biology and Evolution, 16(4), evae062. 10.1093/gbe/evae062

21. Glotova, A., & Smirnov, A. (2017). Description of *Flamella daurica* n. Sp., with notes on the phylogeny of the genus *Flamella* and related taxa. European Journal of Protistology, 58, 164–174. 10.1016/j.ejop.2017.02.003

22. González Miguéns, R. (2023). *Shell morphological diversification patterns and molecular systematics of the testate amoebae orders Arcellinida and Euglyphida* [doctoralThesis]. https://repositorio.uam.es/handle/10486/708114

23. Gordon, D., & Green, P. (2013). Consed: A graphical editor for next-generation sequencing. Bioinformatics, 29(22), 2936–2937. 10.1093/bioinformatics/btt515

24. Grabherr, M. G., Haas, B. J., Yassour, M., Levin, J. Z., Thompson, D. A., Amit, I., Adiconis, X., Fan, L., Raychowdhury, R., Zeng, Q., Chen, Z., Mauceli, E., Hacohen, N., Gnirke, A., Rhind, N., di Palma, F., Birren, B. W., Nusbaum, C., Lindblad-Toh, K., … Regev, A. (2011). Full-length transcriptome assembly from RNA-Seq data without a reference genome. Nature Biotechnology, 29(7), 644–652. 10.1038/nbt.1883

25. Gruber-Vodicka, H. R., Seah, B. K. B., & Pruesse, E. (2020). phyloFlash: Rapid Small-Subunit rRNA Profiling and Targeted Assembly from Metagenomes. mSystems, 5(5), e00920–20. 10.1128/mSystems.00920-20

26. Guéguen, L., Gaillard, S., Boussau, B., Gouy, M., Groussin, M., Rochette, N. C., Bigot, T., Fournier, D., Pouyet, F., Cahais, V., Bernard, A., Scornavacca, C., Nabholz, B., Haudry, A., Dachary, L., Galtier, N., Belkhir, K., & Dutheil, J. Y. (2013). Bio++: Efficient Extensible Libraries and Tools for Computational Molecular Evolution. Molecular Biology and Evolution, 30(8), 1745–1750. 10.1093/molbev/mst097

27. Guerrero, E., Abelló, P., Lombarte, A., Villanueva, R., Ramón, M., Sabatés, A., & Santos, R. (2023). Marine Biological Reference Collections ICM-CSIC. Version 1.30. Instituto de Ciencias del Mar (CSIC). Occurrence dataset 10.15470/qlqqdx [Dataset].

28. Hess, S., Sausen, N., & Melkonian, M. (2012). Shedding Light on Vampires: The Phylogeny of Vampyrellid Amoebae Revisited. PLOS ONE, 7(2), e31165. 10.1371/journal.pone.0031165

29. Hess, S., & Suthaus, A. (2022). The Vampyrellid Amoebae (Vampyrellida, Rhizaria). Protist, 173(1), 125854. 10.1016/j.protis.2021.125854

30. Hülsmann, N., & Grębecki, A. (1995). Induction of lobopodia and lamellipodia in a filopodial organism (*Vampyrella lateritia*). European Journal of Protistology, 31(2), 182–189. 10.1016/S0932-4739(11)80442-6

31. Jeleń, F., Oleksy, A., Smietana, K., & Otlewski, J. (2003). PDZ domains—Common players in the cell signaling. Acta Biochimica Polonica, 50, 985–1017. 10.18388/abp.2003_3628

32. Lê, S., Josse, J., & Husson, F. (2008). FactoMineR: An R Package for Multivariate Analysis. Journal of Statistical Software, 25, 1–18. 10.18637/jss.v025.i01

33. López-García, P., Philippe, H., Gail, F., & Moreira, D. (2003). Autochthonous eukaryotic diversity in hydrothermal sediment and experimental microcolonizers at the Mid-Atlantic Ridge. Proceedings of the National Academy of Sciences, 100(2), 697–702. 10.1073/pnas.0235779100

34. Merényi, Z., Krizsán, K., Sahu, N., Liu, X.-B., Bálint, B., Stajich, J. E., Spatafora, J. W., & Nagy, L. G. (2023). Genomes of fungi and relatives reveal delayed loss of ancestral gene families and evolution of key fungal traits. Nature Ecology & Evolution, 7(8), 1221–1231. 10.1038/s41559-023-02095-9

35. Nguyen, L.-T., Schmidt, H. A., von Haeseler, A., & Minh, B. Q. (2015). IQ-TREE: A Fast and Effective Stochastic Algorithm for Estimating Maximum-Likelihood Phylogenies. Molecular Biology and Evolution, 32(1), 268–274. 10.1093/molbev/msu300

36. Ocaña-Pallarès, E., Williams, T. A., López-Escardó, D., Arroyo, A. S., Pathmanathan, J. S., Bapteste, E., Tikhonenkov, D. V., Keeling, P. J., Szöllősi, G. J., & Ruiz-Trillo, I. (2022). Divergent genomic trajectories predate the origin of animals and fungi. Nature, 609(7928), 747–753. 10.1038/s41586-022-05110-4

37. Park, M. D., Silvin, A., Ginhoux, F., & Merad, M. (2022). Macrophages in health and disease. Cell, 185(23), 4259–4279. 10.1016/j.cell.2022.10.007

38. Pawlowski, J., & Burki, F. (2009). Untangling the Phylogeny of Amoeboid Protists1. Journal of Eukaryotic Microbiology, 56(1), 16–25. 10.1111/j.1550-7408.2008.00379.x

39. Pomorski, P., Krzemiński, P., Wasik, A., Wierzbicka, K., Barańska, J., & Kłopocka, W. (2007). Actin dynamics in Amoeba proteus motility. Protoplasma, 231(1), 31–41. 10.1007/s00709-007-0243-1

40. Raivo Kolde. (2010). *pheatmap: Pretty Heatmaps* (p. 1.0.12) [Dataset]. 10.32614/CRAN.package.pheatmap

41. Richter, D. J., Fozouni, P., Eisen, M. B., & King, N. (2018). Gene family innovation, conservation and loss on the animal stem lineage. eLife, 7, e34226. 10.7554/eLife.34226

42. Ros-Rocher, N., Pérez-Posada, A., Leger, M. M., & Ruiz-Trillo, I. (2021). The origin of animals: An ancestral reconstruction of the unicellular-to-multicellular transition. Open Biology, 11(2), 200359. 10.1098/rsob.200359

43. RStudio Team. (2020). RStudio: Integrated Development for R. *RStudio, PBC, Boston, MA*. http://www.rstudio.com/

44. Schilde, C., & Schaap, P. (2013). The Amoebozoa. In L. Eichinger & F. Rivero (Eds.), *Dictyostelium discoideum Protocols* (pp. 1–15). Humana Press. 10.1007/978-1-62703-302-2_1

45. Schindelin, J., Arganda-Carreras, I., Frise, E., Kaynig, V., Longair, M., Pietzsch, T., Preibisch, S., Rueden, C., Saalfeld, S., Schmid, B., Tinevez, J.-Y., White, D. J., Hartenstein, V., Eliceiri, K., Tomancak, P., & Cardona, A. (2012). Fiji: An open-source platform for biological-image analysis. Nature Methods, 9(7), 676–682. 10.1038/nmeth.2019

46. Sheng, M., & Sala, C. (2001). PDZ domains and the organization of supramolecular complexes. Annual Review of Neuroscience, 24, 1–29. 10.1146/annurev.neuro.24.1.1

47. Siddiqui, R., Ali, I. K. M., Cope, J. R., & Khan, N. A. (2016). Biology and pathogenesis of *Naegleria fowleri*. Acta Tropica, 164, 375–394. 10.1016/j.actatropica.2016.09.009

48. Simão, F. A., Waterhouse, R. M., Ioannidis, P., Kriventseva, E. V., & Zdobnov, E. M. (2015). BUSCO: Assessing genome assembly and annotation completeness with single-copy orthologs. Bioinformatics, 31(19), 3210–3212. 10.1093/bioinformatics/btv351

49. Stensvold, C. R., Ascuña-Durand, K., Chihi, A., Belkessa, S., Kurt, Ö., El-Badry, A., van der Giezen, M., & Clark, C. G. (2023). Further insight into the genetic diversity of Entamoeba coli and Entamoeba hartmanni. Journal of Eukaryotic Microbiology, 70(2), e12949. 10.1111/jeu.12949

50. Suga, H., Chen, Z., de Mendoza, A., Sebé-Pedrós, A., Brown, M. W., Kramer, E., Carr, M., Kerner, P., Vervoort, M., Sánchez-Pons, N., Torruella, G., Derelle, R., Manning, G., Lang, B. F., Russ, C., Haas, B. J., Roger, A. J., Nusbaum, C., & Ruiz-Trillo, I. (2013). The Capsaspora genome reveals a complex unicellular prehistory of animals. Nature Communications, 4, 2325. 10.1038/ncomms3325

51. Tekle, Y. I., Wang, F., Wood, F. C., Anderson, O. R., & Smirnov, A. (2022). New insights on the evolutionary relationships between the major lineages of Amoebozoa. Scientific Reports, 12(1), 11173. 10.1038/s41598-022-15372-7

52. Tice, A. K., Žihala, D., Pánek, T., Jones, R. E., Salomaki, E. D., Nenarokov, S., Burki, F., Eliáš, M., Eme, L., Roger, A. J., Rokas, A., Shen, X.-X., Strassert, J. F. H., Kolísko, M., & Brown, M. W. (2021). PhyloFisher: A phylogenomic package for resolving eukaryotic relationships. PLOS Biology, 19(8), e3001365. 10.1371/journal.pbio.3001365

53. Vaulot, D., Geisen, S., Mahé, F., & Bass, D. (2022). pr2-primers: An 18S rRNA primer database for protists. Molecular Ecology Resources, 22(1), 168–179. 10.1111/1755-0998.13465

54. Wang, Y., Jiang, L., Zhao, Y., Ju, X., Wang, L., Jin, L., Fine, R. D., & Li, M. (2023). Biological characteristics and pathogenicity of Acanthamoeba. Frontiers in Microbiology, 14. 10.3389/fmicb.2023.1147077

55. Wickham, H. (2011). Ggplot2. WIREs Computational Statistics, 3(2), 180–185. 10.1002/wics.147

56. Wuyts, J., De Rijk, P., Van de Peer, Y., Pison, G., Rousseeuw, P., & De Wachter, R. (2000). Comparative analysis of more than 3000 sequences reveals the existence of two pseudoknots in area V4 of eukaryotic small subunit ribosomal RNA. Nucleic Acids Research, 28(23), 4698–4708. 10.1093/nar/28.23.4698

