## Supplemental Figures for "The evolution of gene functional repertoire in Amorphea: Divergent strategies across Amoebozoa, Fungi and Metazoa"

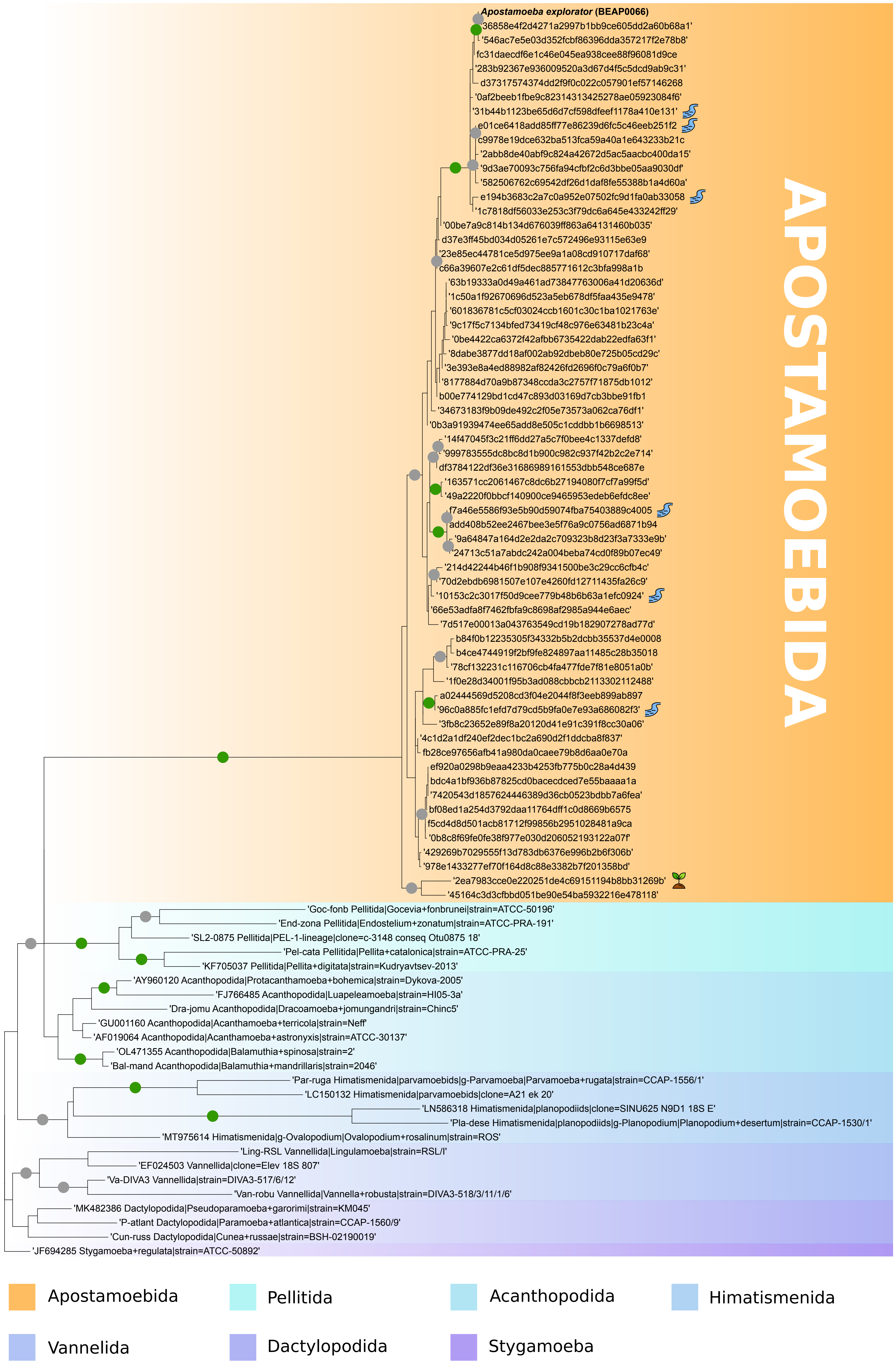


**Supplementary Figure 1. Apostamoebida environmental V4 diversity.** Phylogenetic tree of Apostamoebida diversity based on V4 18S region (330 positions). Phylogeny reconstructed with RAxML using the GTRCAT model of sequence evolution with 1000 rapid bootstrap inferences. Green dots represent a node support over 95, while grey dots represent a node support over 75. Support for the node separating Apostamoebida from other groups is 100. All Apostamoebida representatives, except '2ea7983cce0e220251de4c69151194b8bb31269b', are found in samples taken from a marine environment. Representatives marked with a river symbol or a soil symbol indicate the representative's presence in freshwater and soil samples, respectively.

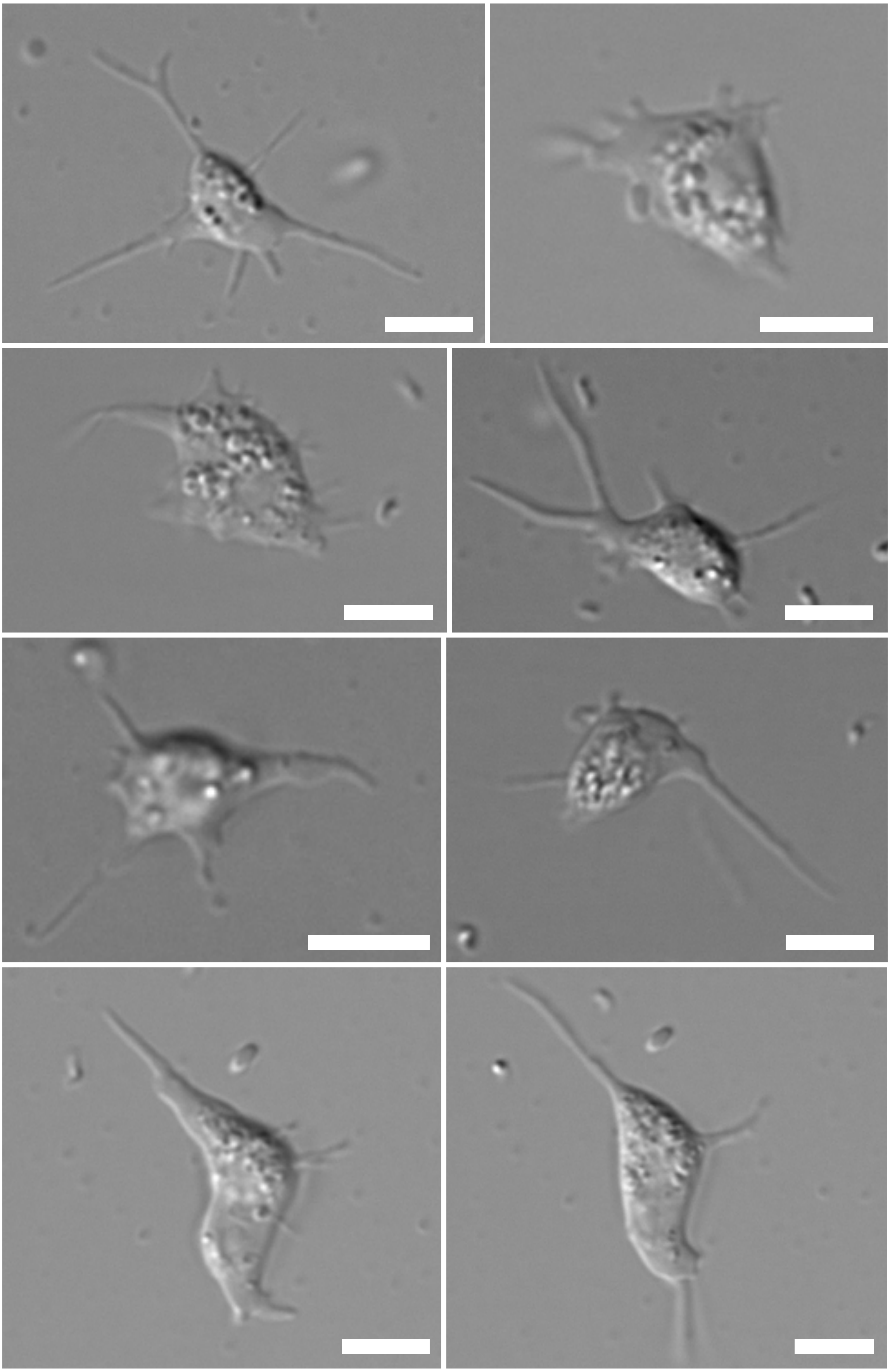

**Supplementary Figure 2. Morphology of *Apostamoeba explorator* through Differential interference contrast (DIC) images.** Differential interference contrast (DIC) images of *Apostamoeba explorator* trophozoites. Scale bars in all panels = 5 µm.

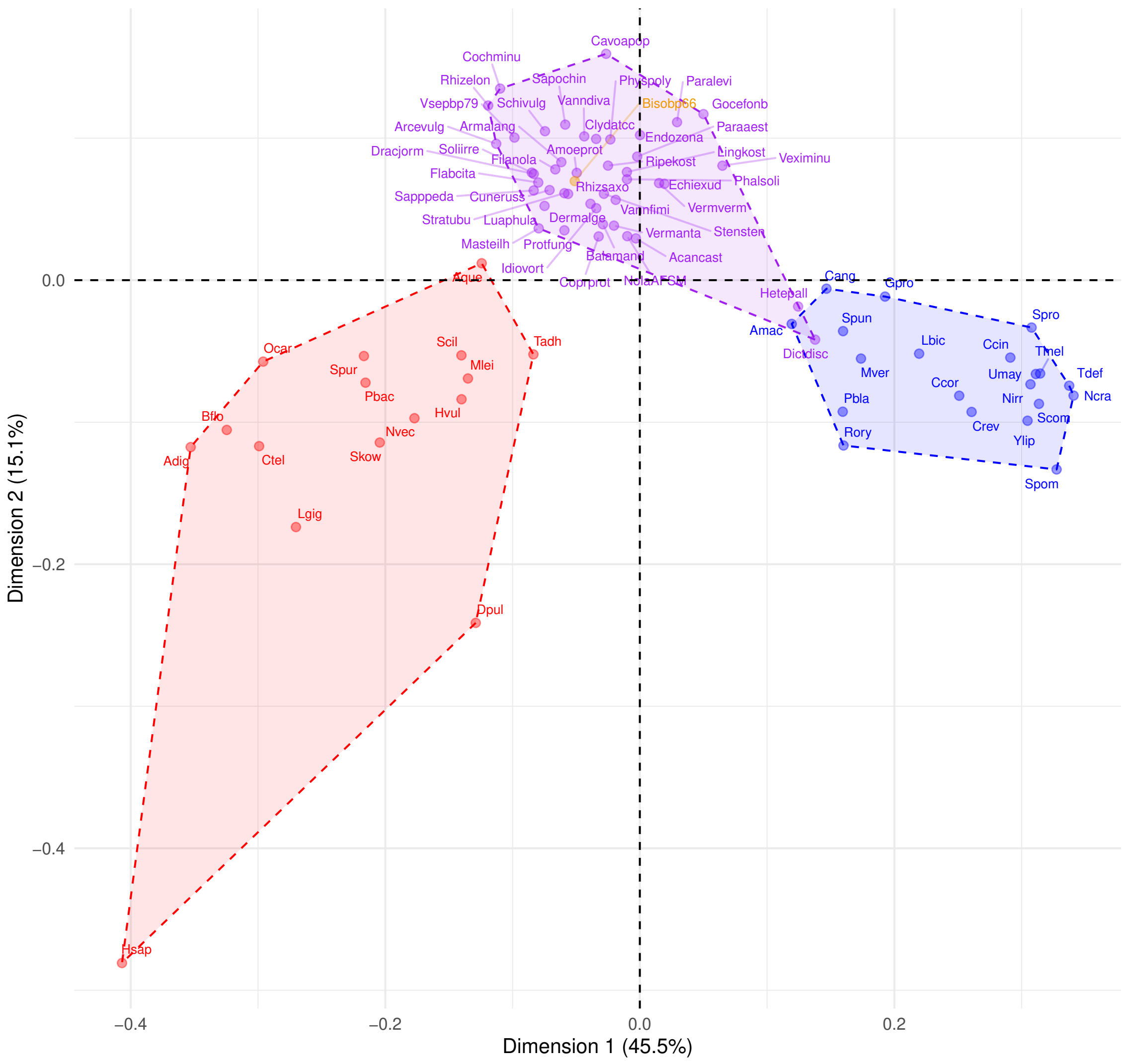


**Taxonomic group**

- Amoebozoa
- Ancestor−Notung
- Fungi
- Metazoa

**COG categories**

A = RNA processing and modification
B = Chromatin structure and dynamics
C = Energy production and conversion
D = Cell cycle control, cell division, chromosome partitioning
E = Amino acid transport and metabolism
F = Nucleotide transport and metabolism
G = Carbohydrate transport and metabolism
H = Coenzyme transport and metabolism
I = Lipid transport and metabolism
J = Translation, ribosomal structure, and biogenesis
K = Transcription
L = Replication, recombination, and repair
M = Cell wall/membrane/envelope biogenesis
N = Cell motility
O = Posttranslational modification, protein turnover, chaperones
P = Inorganic ion transport and metabolism
Q = Secondary metabolites biosynthesis, transport, and catabolism
R = General function prediction only
S = Function unknown
T = Signal transduction mechanisms
U = Intracellular trafficking, secretion, and vesicular transport
V = Defense mechanisms
W = Extracellular structures
Y = Nuclear structure
Z = Cytoskeleton

**Supplementary Figure 3. Correspondence Analyses (CA) of Clusters of Orthologous Groups (COG) functional category compositions of the gene complements of Amoebozoa (purple), Metazoa (red) and Fungi (blue) with all species labeled.** First two dimensions of CA. Each point represents the gene complement of a single species (from a genome or a transcriptome; Supplementary figure 5B shows that data source does not have a significant impact). Correspondence between the species abbreviations and their full names can be found in Supplementary File 2.

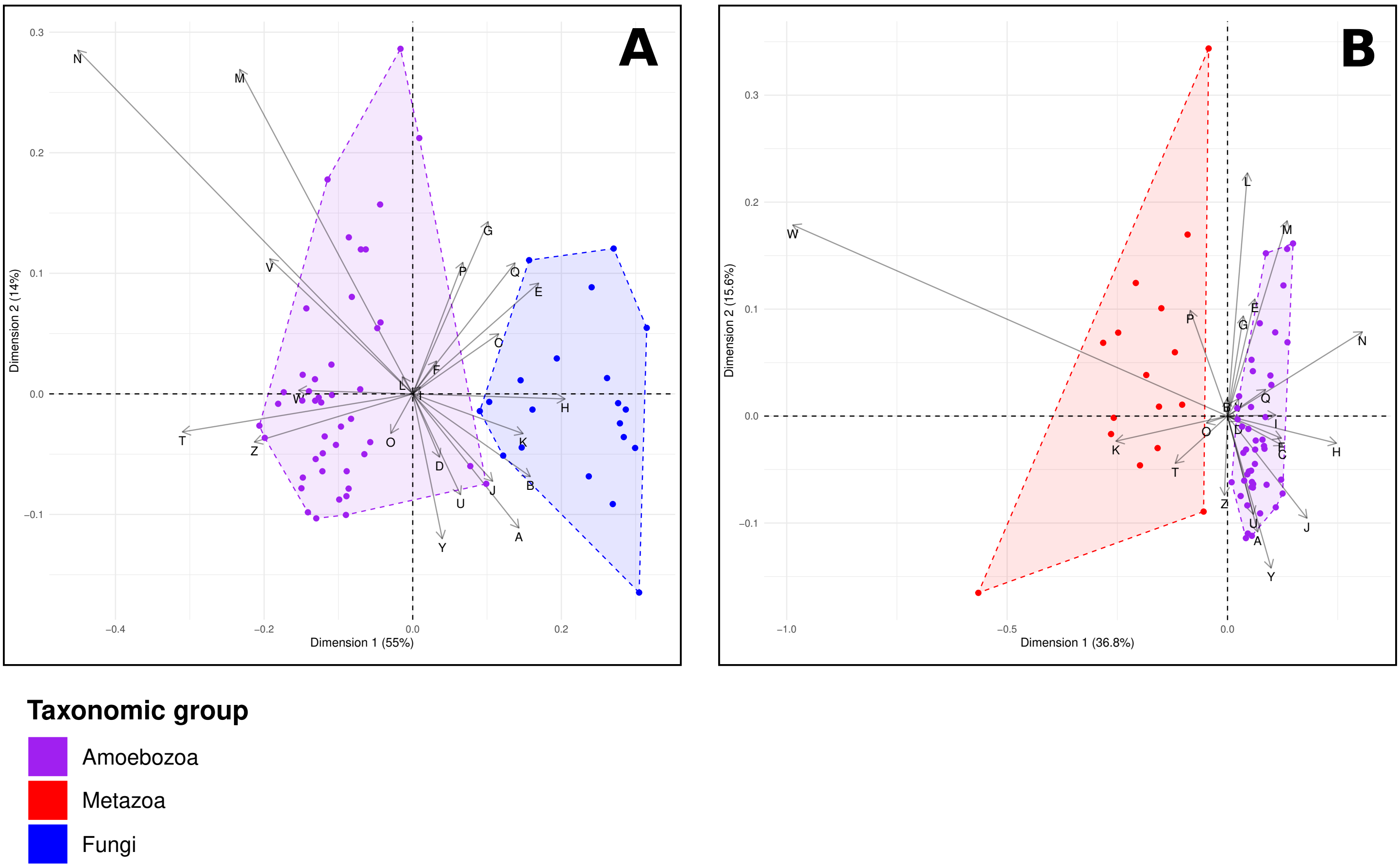


**Supplementary Figure 4. Correspondence Analyses (CA) of Clusters of Orthologous Groups (COG) functional category compositions of the gene complements of Amoebozoa against either Fungi or Metazoa.** First two dimensions of CA. Each point represents the gene complement of a single species (from a genome or a transcriptome; Supplementary figure 5B shows that data source does not have a significant impact). **A.** Amoebozoa (purple) compared with Fungi (Blue). **B.** Amoebozoa (Purple) compared with Metazoa (Red).

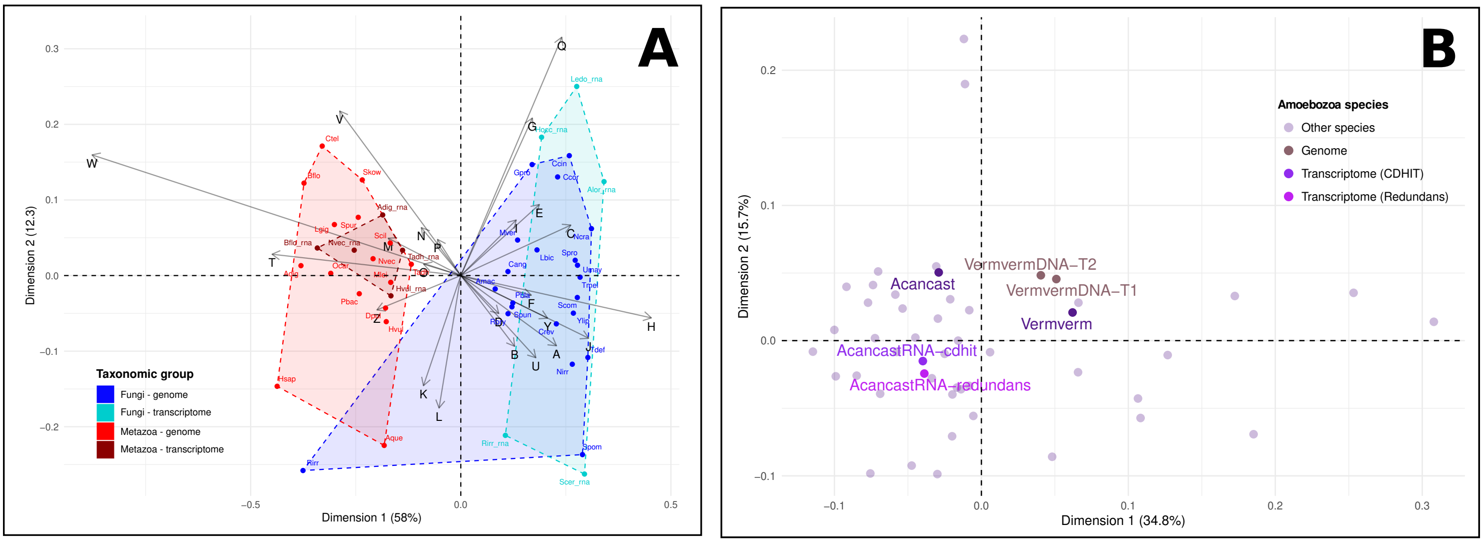


**Supplementary Figure 5. Correspondence Analyses (CA) of Clusters of Orthologous Groups (COG) functional category compositions of gene complements estimated with different methods. A.** Metazoa (red) and Fungi (blue) genomic and transcriptomic gene complements are compared and cluster together. Five transcriptomes per taxonomic group were selected, each with a genomic counterpart from our study (same species or closely related). All used datasets are publicly available. The genome complement of Rirr (*Rhizopus irregularis*), which was excluded from our other analysis, does not cluster with other fungi, although its transcriptome complement is less of an outlier. **B.** Amoebozoa genomic (brown) and transcriptomic (purple) gene complements compared with their counterparts. *Acanthamoeba castellani* (Acancast) transcriptome was processed separately with either CD-HIT or Redundans, in order to reduce redundancy and to assess the equivalence of these programs, and compared with its genome (originally used in the project). Genes in the *Vermamoeba vermiformis* (Vermverm) genome were predicted with EukMetaSanity and compared with its transcriptome (originally used in the project).

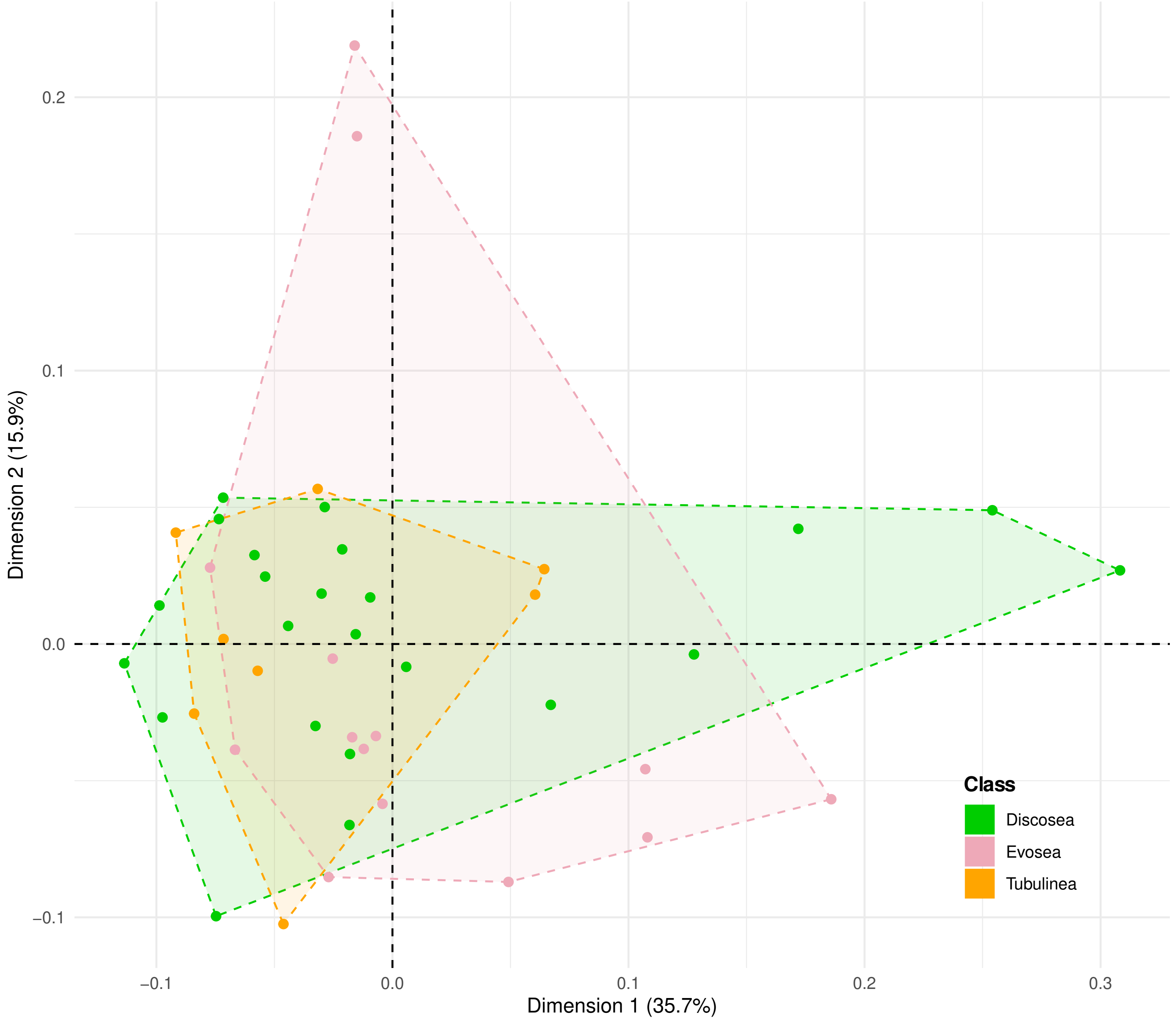


**Supplementary Figure 6. Correspondence Analyses (CA) of Clusters of Orthologous Groups (COG) functional category compositions of the gene complements of groups within Amoebozoa.** Discosea (green), Tubulinea (orange) and Evosea (pink). First two dimensions of CA. Each point represents the gene complement of a single species (from a genome or a transcriptome; Supplementary figure 5B shows that data source does not have a significant impact).

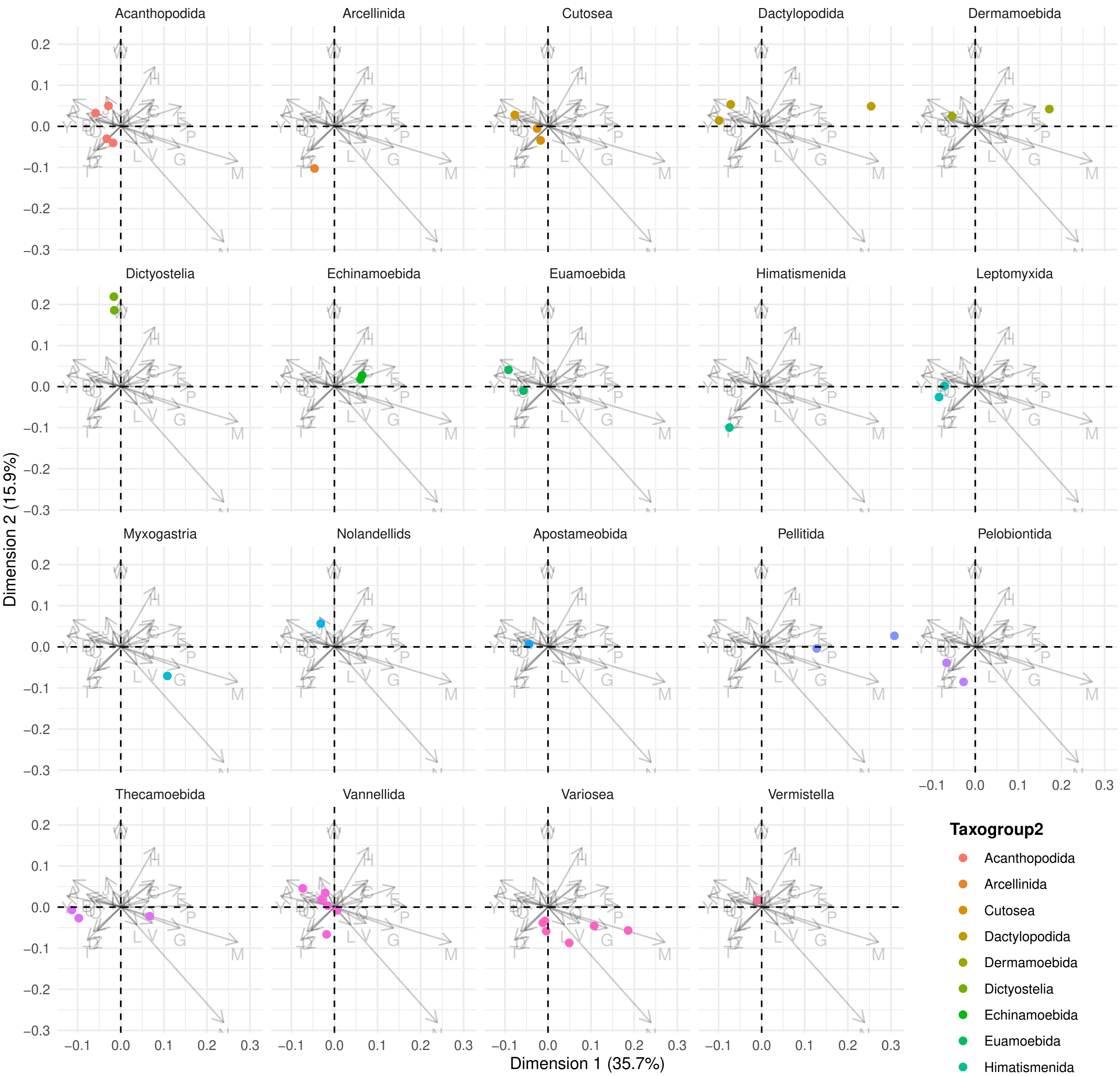


**Supplementary Figure 7. Correspondence Analyses (CA) of Clusters of Orthologous Groups (COG) functional category compositions of the gene complements displayed for individual groups within Amoebozoa.** Taxonomic groupings are taken from UniEuk, at the Taxogroup2 level (Berney et al., 2017). A single CA was performed. Different panels highlight members of each Taxogroup2, in color. First two dimensions of CA. Each point represents the gene complement of a single species (from a genome or a transcriptome; Supplementary figure 5B shows that data source does not have a significant impact).

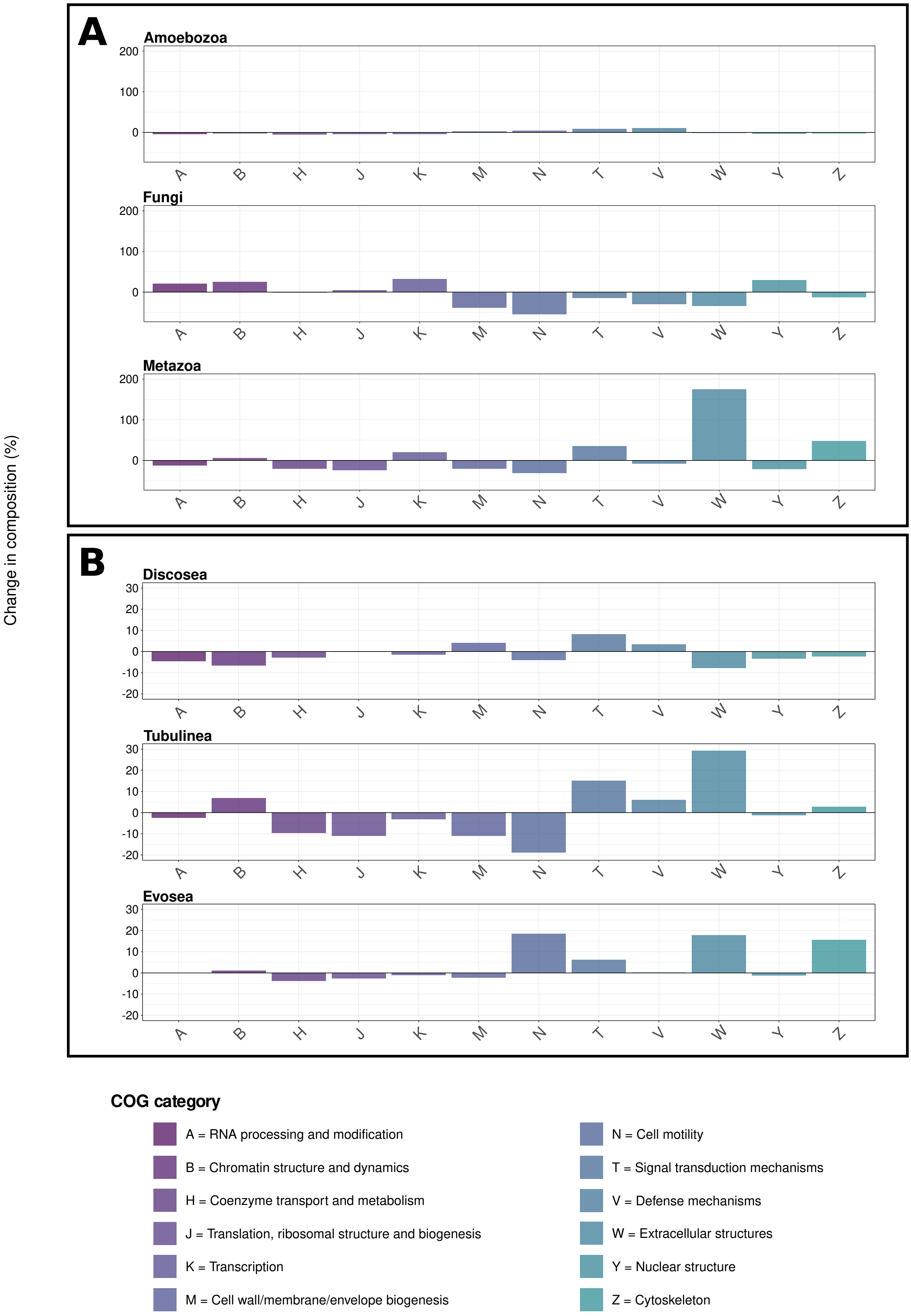


**Supplementary Figure 8. Reconstructed ancestral gains and losses in Clusters of orthologous groups (COG) functional category composition. A.** Changes on the stem lineage leading from Amorphea to the last common ancestors of Amoebozoa, Fungi or Metazoa. **B.** Changes on the stem lineage leading from Amoebozoa to the last common ancestors of Discosea, Tubulina or Evosea. Only a subset of COG categories are represented in the figure, specifically those with a change greater than 20% from the Amorphea ancestor to either Amoebozoa, Fungi, or Metazoa.

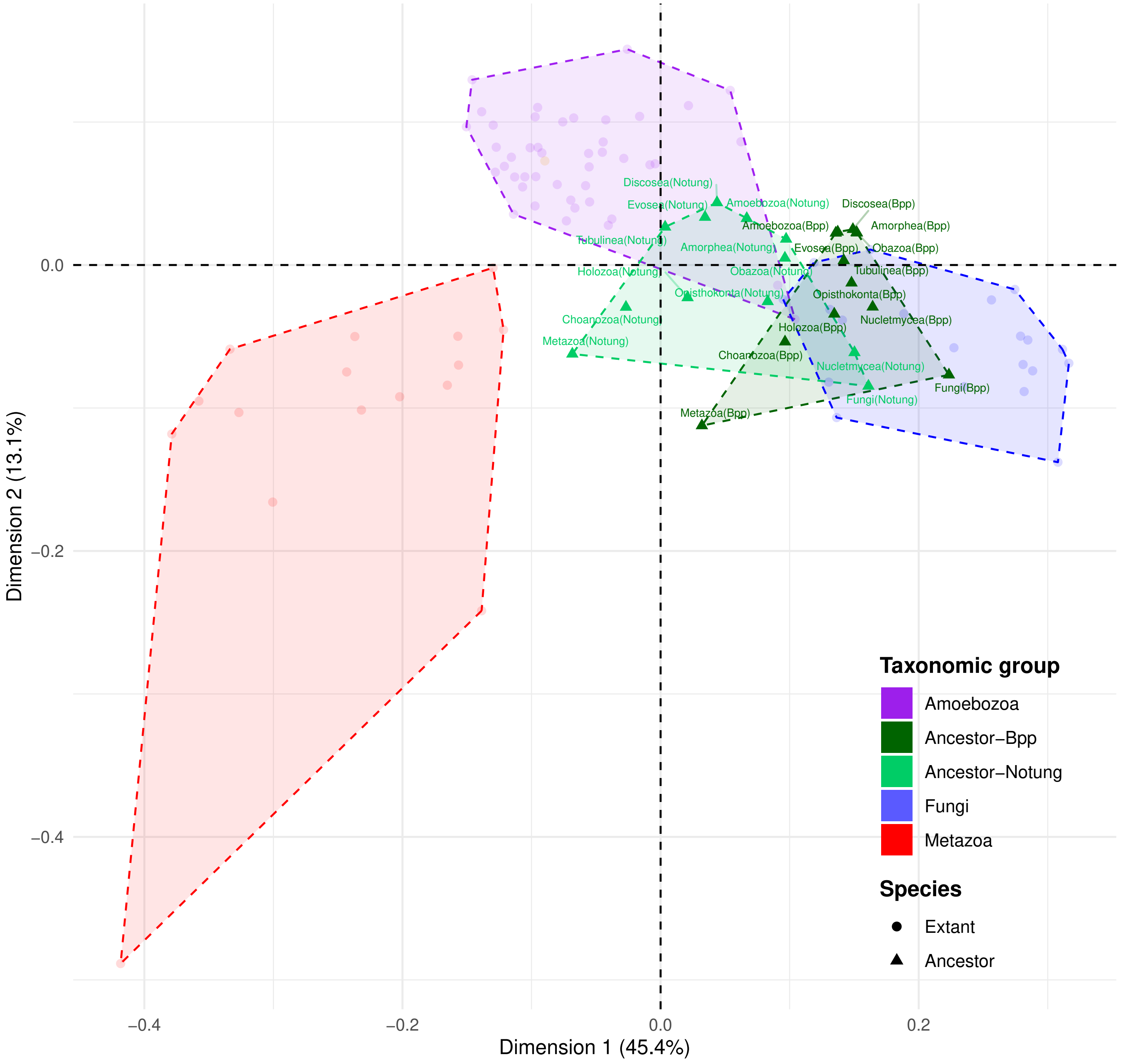


**Supplementary Figure 9. Correspondence Analyses (CA) of Clusters of Orthologous Groups (COG) functional category compositions of the gene complements of Amoebozoa, Metazoa, Fungi and their most recent common ancestors.** First two dimensions of CA. Each point represents the gene complement of a single species (from a genome or a transcriptome; Supplementary figure 5B shows that data source does not have a significant impact). Gene complements of most recent common ancestors were inferred by orthologous gene identification (OrthoFinder2) and either Wagner parsimony (Notung) or maximum likelihood (Bpp). Wagner parsimony estimates ancestral counts for each OrthoFinder2 gene family, whereas maximum likelihood with Bpp estimates ancestral presence or absence.

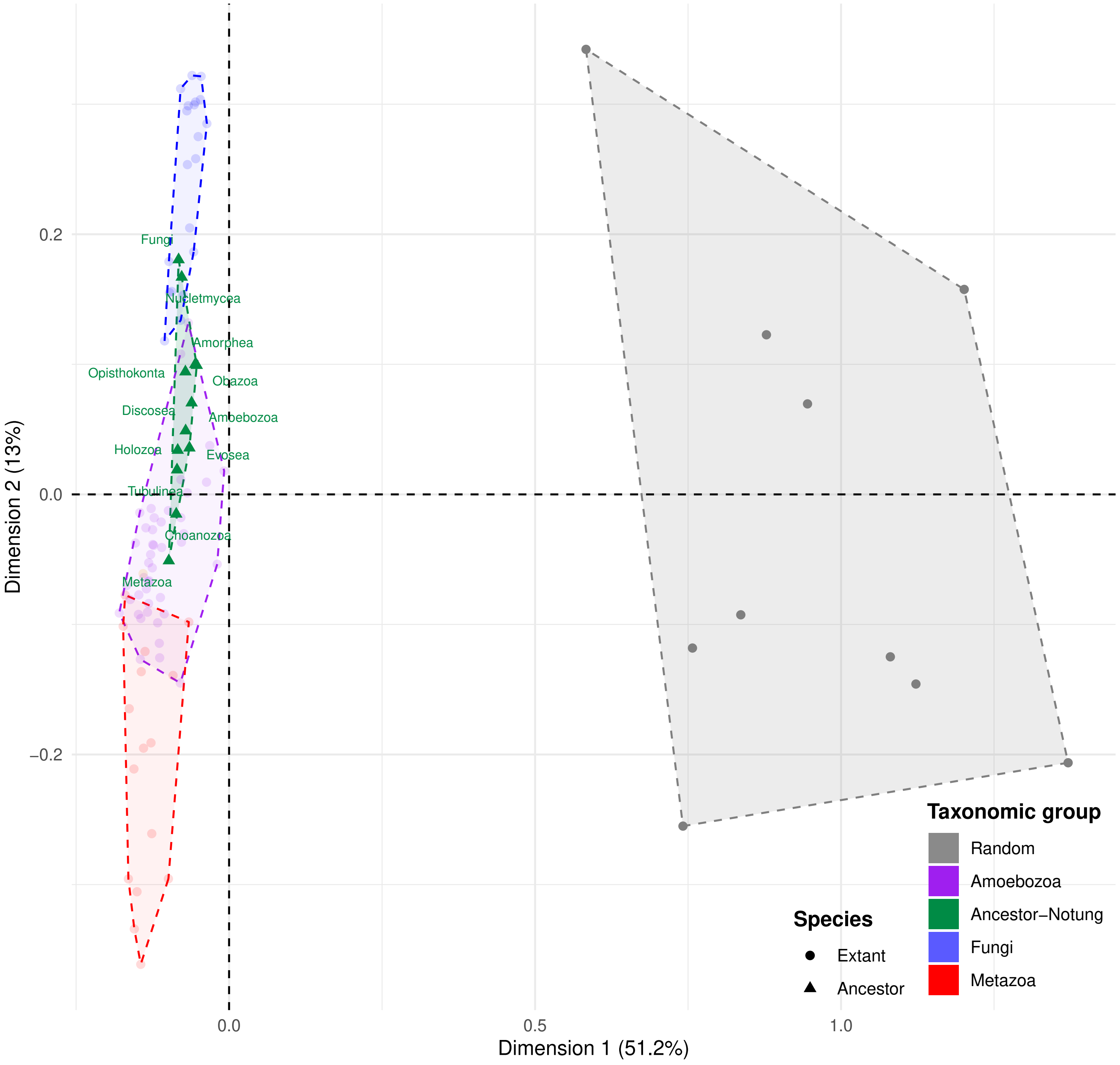


**Supplementary Figure 10. Correspondence Analyses (CA) of Clusters of Orthologous Groups (COG) functional category compositions of the gene complements of Amoebozoa (purple), Metazoa (red), Fungi (blue) and their most recent common ancestors (green), together with randomly generated gene complements (grey).** First two dimensions of CA. Each point represents the gene complement of a single species (from a genome or a transcriptome; Supplementary figure 5B shows that data source does not have a significant impact). Most recent common ancestors with gene complements were inferred by orthologous gene identification (OrthoFinder2) followed by Wagner parsimony (Notung). The points in the different random sets clusters are distant from the species we analyzed.
